# Context-dependent requirement of G protein coupling for Latrophilin-2 in target selection of hippocampal axons

**DOI:** 10.1101/2022.09.26.509559

**Authors:** Daniel T. Pederick, Nicole A. Perry-Hauser, Huyan Meng, Zhigang He, Jonathan A. Javitch, Liqun Luo

## Abstract

The formation of neural circuits requires extensive interactions of cell-surface proteins to guide axons to their correct target neurons. *Trans*-cellular interactions of the adhesion G protein-coupled receptor latrophilin-2 (Lphn2) with its partner teneurin-3 instruct the precise assembly of hippocampal networks by reciprocal repulsion. Lphn2 acts as a receptor in distal CA1 neurons to direct their axons to proximal subiculum, and as a repulsive ligand in proximal subiculum to direct proximal CA1 axons to distal subiculum. It remains unclear if Lphn2-mediated intracellular signaling is required for its role in either context. Here, we show that Lphn2 couples to Gα_12/13_ in heterologous cells, which is increased by constitutive exposure of the tethered agonist. Specific mutations of Lphn2’s tethered agonist region disrupt its G protein coupling and autoproteolytic cleavage, whereas mutating the autoproteolytic cleavage site prevents cleavage but preserves a functional tethered agonist. Using an *in vivo* misexpression assay, we demonstrate that wild-type Lphn2 misdirects proximal CA1 axons to proximal subiculum and that Lphn2 tethered agonist activity is required for its role as a repulsive receptor. By contrast, neither tethered agonist activity nor autoproteolysis was necessary for Lphn2’s role as a repulsive ligand. Thus, tethered agonist activity is required for Lphn2-mediated neural circuit assembly in a context-dependent manner.

## INTRODUCTION

Latrophilins (Lphn1-3) are highly expressed in the brain and were originally identified as responders to ɑ-latrotoxin, a neurotoxin from black widow spider venom that causes profound release of neurotransmitters from nerve terminals (Davletov et al., 1996). They belong to the family of adhesion G protein-coupled receptors (aGPCRs), capable of eliciting intracellular effects through coupling with heterotrimeric G proteins (Lelianova et al., 1997). Additionally, as cell adhesion molecules, latrophilins interact via their N-terminal extracellular domain with four different families of interacting partners including neurexins (Boucard et al., 2012), teneurins (Silva et al., 2011), fibronectin leucine-rich transmembrane proteins (FLRTs) (O’Sullivan et al., 2012), and contactins (Zuko et al., 2016). In the central nervous system, latrophilins have been implicated in neuronal migration, circuit assembly, and synapse formation (Anderson et al., 2017; Del Toro et al., 2020; Donohue et al., 2021; Pederick et al., 2021; Sando and Südhof, 2021; Sando et al., 2019). In humans, mutation of *LPHN3* is associated with increased risk of attention deficit/hyperactivity disorder and a missense variant in the *LPHN2* gene is responsible for extreme microcephaly (Arcos-Burgos et al., 2010; Domené et al., 2011; Vezain et al., 2018).

We recently showed in mice that one of the three latrophilins, Lphn2, displays expression inverse to teneurin-3 (Ten3) in two parallel hippocampal networks (Pederick et al., 2021). While hippocampal *Lphn2* is preferentially expressed in the distal CA1 and the proximal subiculum, *Ten3* is enriched in the proximal CA1 and the distal subiculum. These expression patterns and reciprocal repulsions mediated by Ten3-Lphn2 interactions instruct proximal CA1 axons to target distal subiculum, and more distal CA1 axons to target more proximal subiculum (**Figure 1A**). Specifically, Lphn2 acts as a “receptor” in more distal CA1 axons that is repelled by Ten3 expressed from distal subiculum (**Figure 1B**). At the same time, Lphn2 acts as a repulsive “ligand” in the proximal subiculum to repel Ten3+ proximal CA1 axons; this action requires Lphn2’s teneurin-binding domain but not its FLRT-binding activity (**Figure 1C**) (Pederick et al., 2021). Therefore, Lphn2 is required cell autonomously as a receptor in more distal CA1 axons for their precise target selection, and non-autonomously in target neurons as a ligand for precise target selection of proximal CA1 axons.

**Figure 1.**
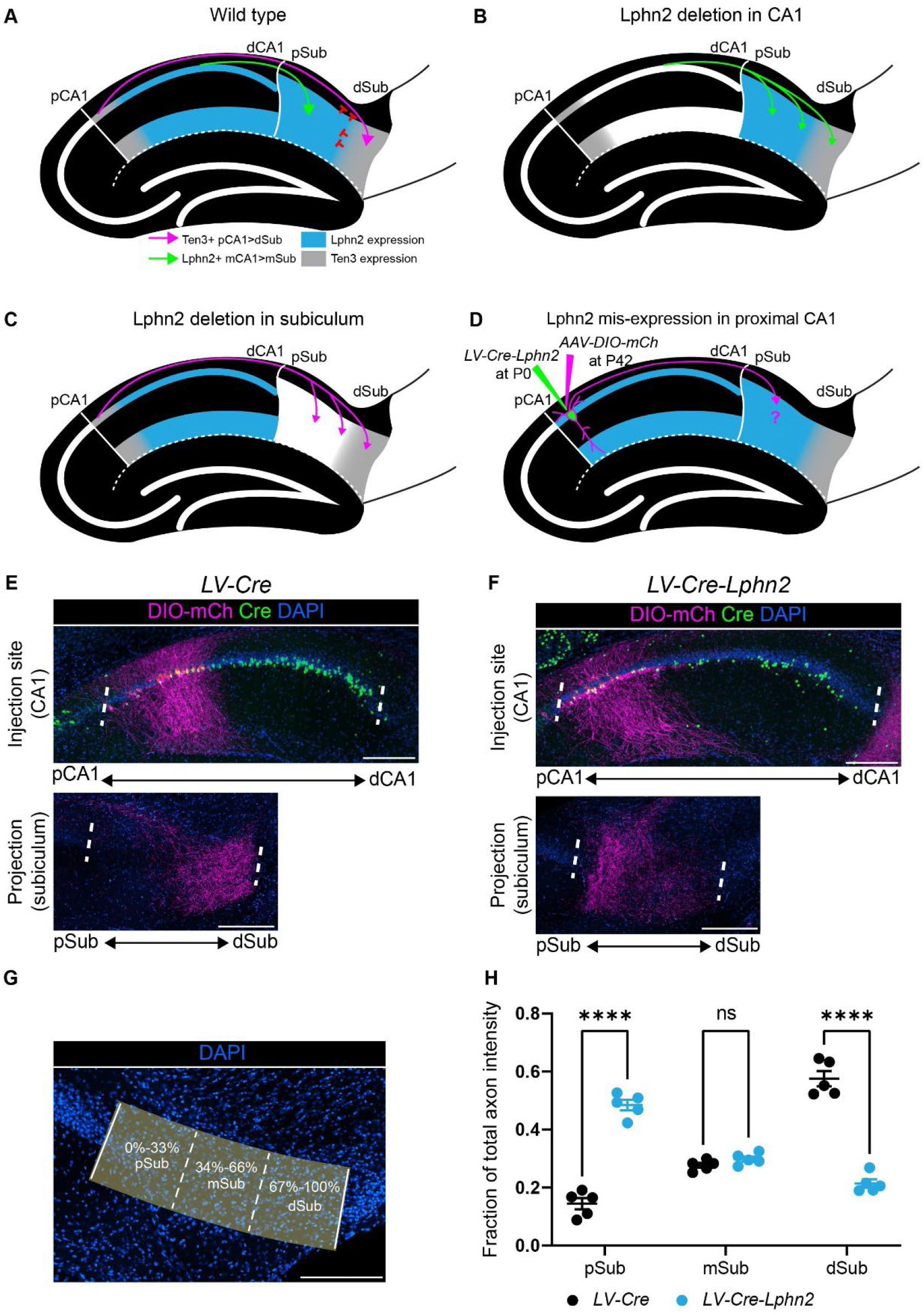
Misexpression of Lphn2 in proximal CA1 axons causes axon mistargeting to the proximal subiculum. **(A)** Cartoon depicting the topographic connections from proximal CA1 (pCA1) to distal subiculum (dSub) and distal CA1 (dCA1) to proximal subiculum (pSub). Ten3+ proximal CA1 axons are repelled from Lphn2+ proximal subiculum and Lphn2+ axons are repelled from Ten3+ distal subiculum. **(B)** Deletion of Lphn2 from CA1 leads to distal CA1 axons mistargeting to distal subiculum, suggesting that Lphn2 acts cell-autonomously as a repulsive receptor. **(C)** Deletion of Lphn2 from proximal subiculum results in proximal CA1 axon mistargeting to proximal subiculum, suggesting Lphn2 acts cell-non-autonomously as a repulsive ligand. Figures (A-C) are based on (Pederick et al., 2021). **(D)** Experimental design of Lphn2 misexpression assay in proximal CA1. At postnatal day (P) 0, lentivirus expressing Cre or Cre + Lphn2 was injected into CA1. This was followed by injection at P42 of Cre-dependent membrane bound mCherry (mCh) into proximal CA1 as an axon tracer. **(E** and **F)** Representative images of *AAV-DIO-mCh* (magenta; mCh expression in a Cre-dependent manner) injections in proximal CA1 (top) and corresponding projections in the subiculum (bottom). **(G)** A representative image of the subiculum with proximal subiculum (pSub), mid subiculum (mSub) and distal subiculum (dSub) regions highlighted. **(H)** The fraction of total axon intensity within proximal, mid, and distal subiculum. *LV-Cre*: n=5 and *LV-Cre-Lphn2*: n=5. Means ± SEM; two-way ANOVA with Sidak’s multiple comparisons test. Injection sites of all subjects are shown in **Figure 1—figure supplement 2**. Scale bars represent 200 μm.

Structurally, the N-terminal extracellular domain of latrophilins comprises a rhamnose-binding lectin (RBL) domain, an olfactomedin-like (OLF) ligand-binding domain, a serine/threonine-rich region and hormone receptor motif (HRM), and a conserved GPCR autoproteolysis-inducing (GAIN) domain that encompasses the GPCR proteolysis site (GPS) (Araç et al., 2012; Moreno-Salinas et al., 2019; Vizurraga et al., 2020) (**Figure 2A**). aGPCRs undergo autoproteolytic cleavage at the HL/T consensus site within the GPS. This self-cleavage divides the receptor into an extracellular N-terminal fragment (NTF) and membrane-bound C-terminal fragment (CTF) that remain noncovalently associated throughout biosynthesis and membrane trafficking (Vizurraga et al., 2020). The seven residues immediately C-terminal to the GPS constitute the tethered agonist peptide (TA) (also known as the Stachel or stalk peptide), which upon exposure binds within the transmembrane domain to activate heterotrimeric G proteins (Liebscher and Schöneberg, 2016).

**Figure 2.**
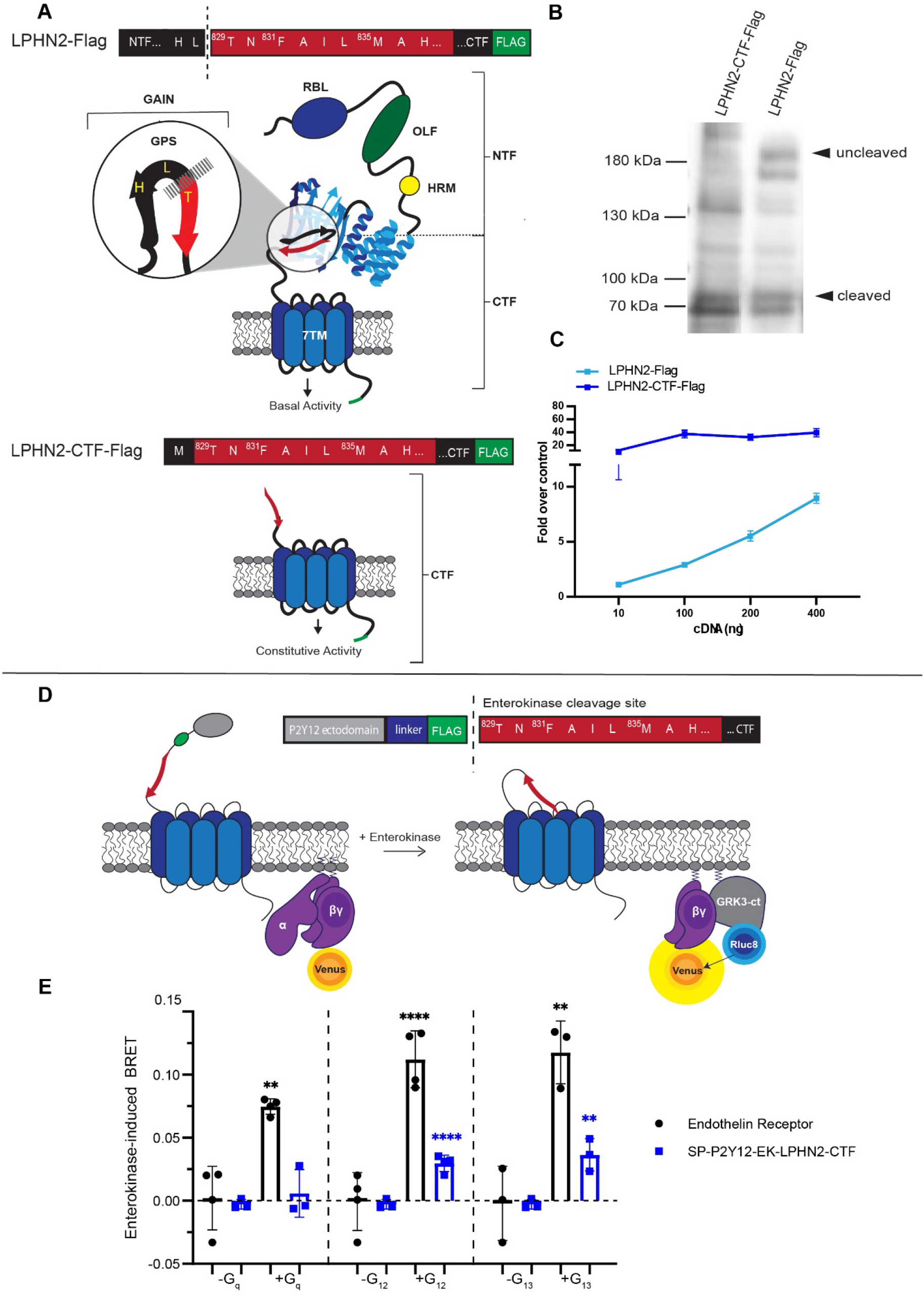
Exposure of the Lphn2 tethered agonist (TA) promotes intracellular signaling through Gα_12/13_. **(A)** Cartoon representations of FL and TA-exposed (CTF) Lphn2 with detailed amino acid sequences for the TA. The extracellular domain of Lphn2 comprises an N-terminal rhamnose-binding lectin domain (RBL), an olfactomedin-like domain (OLF), a serine/threonine-rich region, and a HormR domain. It also contains the GPCR autoproteolysis-inducing (GAIN) domain necessary for autoproteolytic cleavage. This cleavage divides the aGPCR into two polypeptide chains: an N-terminal fragment (NTF) and a C-terminal fragment (CTF). The peptide stretch directly following the proteolytic cleavage site is known as the “Stachel” or tethered agonist (TA). Exposure of the TA results in aGPCR activation and downstream signaling. **(B)** Immunoblot analysis of WT Lphn2 and Lphn2-CTF expression in HEK293T cells using a primary antibody against Flag (1:500, ThermoFisher, PA1-984B). Expected bands for FL Lphn2-Flag and Lphn2-CTF-Flag are 164 kDa and 72 kDa, respectively. **(C)** SRE luciferase reporter assay for Lphn2 and Lphn2-CTF shows that removing the entire NTF up to the GPS cleavage site constitutively enhances SRE signaling. **(D)** Schematic outlining the Gβγ-release BRET assay. The Lphn2 TA is capped with an enterokinase cleavage site (EK) preceded by a hemagglutinin signal peptide (SP), the P2Y12 N-terminal extracellular sequence, and a flexible linker (Lizano et al., 2021). Addition of 10 nM enterokinase generates a TA neoepitope identical to activated endogenous Lphn2. Lphn2 activation results in G protein dissociation, allowing Gβγ-Venus to associate with the C-terminus of GPCR kinase 3 (GRK3-ct) (Hollins et al., 2009). **(E)** Gβγ-release BRET assay testing SP-P2Y12-EK-Lphn2-CTF activation of Gα_q_, Gα_12_, and Gα_13_ in HEKΔ7 cells. Endothelin receptor with 100 nM ligand ET-1 is used as a positive control. Means ± SEM; Multiple unpaired t tests between no G protein and G protein conditions; **, p<0.01; ****, p<0.0001.

While our previous *in vivo* work established that interaction between Ten3 and Lphn2 was required for precise circuit assembly (Pederick et al., 2021), it did not examine how this might depend on Lphn2-mediated signaling mechanisms. Here, we modified our previous hippocampal model to develop a Lphn2 misexpression assay (**Figure 1D)**. We misexpressed Lphn2 in either CA1 axons or the subiculum target and assessed the impact on normal proximal CA1→distal subiculum axon targeting. We found that ectopically expressing wild-type (WT) Lphn2 in proximal CA1 axons causes their mistargeting to the proximal subiculum. This provided us with a robust platform to interrogate whether TA activity or autoproteolytic cleavage is required for axon targeting in the Lphn2 misexpression system. When misexpressed in CA1, Lphn2 TA activity was required for Lphn2-mediated axon targeting. By contrast, when we misexpressed Lphn2 in subiculum target neurons, both TA activity and autoproteolysis were dispensable for Lphn2-mediated axon repulsion. Thus, our data support that Lphn2 G protein coupling is required in axons but not target neurons during precise circuit assembly.

## RESULTS

### Misexpression of WT Lphn2 in proximal CA1 leads to axon mistargeting in the subiculum

To investigate the role of Lphn2-mediated G protein activity in hippocampal axon targeting, we first designed a gain-of-function assay in which we misexpressed Lphn2 in proximal CA1 neurons. We hypothesized that this ectopic expression would cause proximal CA1 axons to avoid Ten3-positive distal subiculum and incorrectly target the proximal subiculum, and that this platform could provide us an assay to test Lphn2 mutants with defects in various functions to determine whether WT Lphn2 mistargeting is compromised.

To test our hypothesis, we used a dual injection strategy to ectopically express Lphn2 in proximal CA1 and trace its axons into the subiculum (**Figure 1—figure supplement 1**). At postnatal day 0 (P0), lentivirus expressing *Cre* (*LV-Cre*) (control) or *Cre-Lphn2* (*LV-Cre-Lphn2*) was injected into proximal CA1, followed by injection of a Cre-dependent membrane-bound mCherry (*AAV-DIO-mCherry*) into proximal CA1 in the same mice at approximately P42 (**Figure 1D**). As expected, in control animals (*LV-Cre*), Cre+ proximal CA1 axons targeted the most distal parts of the subiculum (**Figure 1E**). By contrast, when Lphn2 was misexpressed in proximal CA1 (*LV-Cre-Lphn2*), Cre+ proximal CA1 axons targeted the most proximal parts of the subiculum (**Figure 1F**). To analyze the location of proximal CA1 axons in the subiculum, we calculated the fraction of axon intensity within thirds of the subiculum across the proximal/distal axis (**Figure 1G**). Proximal CA1 axons misexpressing Lphn2 are located significantly more in the proximal third of the subiculum and significantly less in the distal third of the subiculum when compared to control axons (**Figure 1H**).

These data supported our hypothesis that ectopic expression of Lphn2 in proximal CA1 axons causes mistargeting to the proximal subiculum. Having established the effect of wild-type (WT) Lphn2 misexpression in proximal CA1 axons, we next sought to characterize G protein coupling of WT Lphn2 and generate Lphn2 mutants to test the requirement of G protein signaling in Lphn2 mediated neural circuit assembly.

### WT Lphn2 signals through Gα_12/13_

The G protein interaction partners for Lphn2 have not been previously established. We recently showed that Lphn3, another member of the latrophilin family of aGPCRs, couples principally to Gα_12/13_, and also more weakly to Gα_q_, using a combination of gene expression assays and an activation strategy that permitted acute exposure of the TA in a live-cell system (Mathiasen et al., 2020). Thus, we began our signaling characterization of Lphn2 similarly using a WT full-length (FL) Lphn2 construct, and a constitutively active construct termed Lphn2-CTF (**Figure 2A**). The WT Lphn2 construct comprises all extracellular elements including the RBL and OLF domains, the HRM, and the GAIN domain, in addition to the 7 transmembrane helix domain. The Lphn2-CTF lacks the entire NTF up to the GPS and instead has only a methionine residue before the TA. We tested expression of these constructs in mammalian cells using immunoblotting and showed that Lphn2-CTF ran at the expected truncated position (∼72 kDa) and that WT Lphn2 ran at both molecular weights corresponding to uncleaved (∼164 kDa) and cleaved (∼72 kDa) positions (**Figure 2B**). This result for FL Lphn2 is similar to previous work that characterized autoproteolysis of Lphn1 (Araç et al., 2012) as well as to our recent findings in Lphn3 (N.A. Perry-Hauser, unpublished data).

To infer activity of these constructs in G protein signaling pathways, we used the luminescence-based gene expression assay for serum response element (SRE), which produces a robust response in our previous studies of Lphn3 (Mathiasen et. al., 2020). In our assay design, SRE action is coupled to the transcription and translation of firefly luciferase; this readout is then normalized to the control reporter, *Renilla* luciferase, expressed from the same plasmid under a constitutive promoter. We found that Lphn2-CTF significantly enhanced signaling over WT Lphn2 for SRE gene expression at varying levels of cDNA transfection (**Figure 2C**). Since the SRE assay reports on signaling by G12/G13 as well as Gq, we tested whether Gα_12/13_ or Gα_q_ was the primary contributor to this response using a selective Gα_q_ inhibitor, YM-254890 (**Figure 2—figure supplement 1**). We did not observe a significant effect upon addition of the inhibitor, suggesting that Lphn2 signals through Gα_12/13_.

To verify our result in the context of acute G protein activation, we next tested how TA exposure affects G protein activation in a bioluminescence resonance energy transfer (BRET) assay (**Figure 2D**). We designed a synthetically-activatable Lphn2 construct based on a recent publication that took advantage of the protease enterokinase (Lizano et al., 2021). Enterokinase selectively recognizes the trypsinogen substrate sequence DDDDK and cleaves after the lysine residue, thereby exposing the native TA. Thus, we cloned a Lphn2 construct that included a modified hemagglutinin signal peptide, the P2Y12 N-terminal extracellular sequence (amino acids 1-24), a flexible linker (GGSGGSGGS), the enterokinase recognition site (DYKDDDDK), and the truncated Lphn2-CTF sequence. We tested this construct in a Gβγ-release assay where energy transfer was monitored between the membrane-anchored luminescent donor, GRK3-ct-Rluc8, and the fluorescent acceptor, Gγ-Venus (Hollins et al., 2009). This assay was performed in a HEKΔ7 cell line with targeted deletion of Gα_12_ and Gα_13,_ as well as Gα_s/olf_, Gα_q/11_, and Gα_z_ (Alvarez-Curto et al., 2016) to enable systematic re-introduction of the Gα subunits. As expected, in the absence of Gα subunits no BRET signal was observed; however, when Gα_12_ or Gα_13_ was re-introduced to cells expressing the Lphn2 construct there was a significant increase in the BRET signal upon treatment with enterokinase (**Figure 2E**). This increase was not observed upon co-expression of the receptor with Gα_s_, Gα_i1,_ or Gα_q_ (**Figure 2—figure supplement 2**). This suggests that the increase in cAMP reported previously for the Lphn2 CTF does not result from direct activation of Gs, but rather from some other form of signaling crosstalk (Sando and Südhof, 2021).

Taken together, these data demonstrate that Lphn2 signals through the G proteins Gα_12_ and Gα_13_. Having established that these in-cell methods were sufficient to characterize G protein signaling pathways for Lphn2, we next characterized how different mutations in the TA region affect intracellular signaling.

### Mutating conserved residues F831A and M835A in the TA impairs G protein-coupling activity

Previous studies suggest that the third and seventh residues of aGPCRs are required for TA-mediated G protein activation (Stoveken et al., 2015). We hypothesized that mutating these residues in Lphn2, phenylalanine (F831) and methionine (M835), to alanine (F831A/M835A) would impair G protein signaling mediated by the TA (**Figure 3A**). Like our work with WT Lphn2 and Lphn2-CTF, we mutated the TA residues in both FL and truncated constructs (Lphn2-F831A/M835A and Lphn2-CTF-F831A/M835A, respectively). Immunoblotting against the C-terminal Flag-tag confirmed expression in HEK293T cells but showed that a substantial portion of Lphn2-F831A/M835A is uncleaved (**Figure 3B**). This is consistent with previous work with Lphn1 showing that mutating the third phenylalanine to an alanine disrupts autoproteolytic cleavage (Araç et al., 2012) and shows that the double mutation (F831A/M835A) in Lphn2 also inhibits cleavage. We also validated that Lphn2-F831A/M835A is expressed on the cell surface at a comparable level as Lphn2 WT (**Figure 3—figure supplement 1)**. We then proceeded to test these constructs in our SRE gene expression system (**Figure 3A**). As hypothesized, both the FL and truncated Lphn2 had dramatically impaired responses to SRE across varying levels of cDNA transfection.

**Figure 3.**
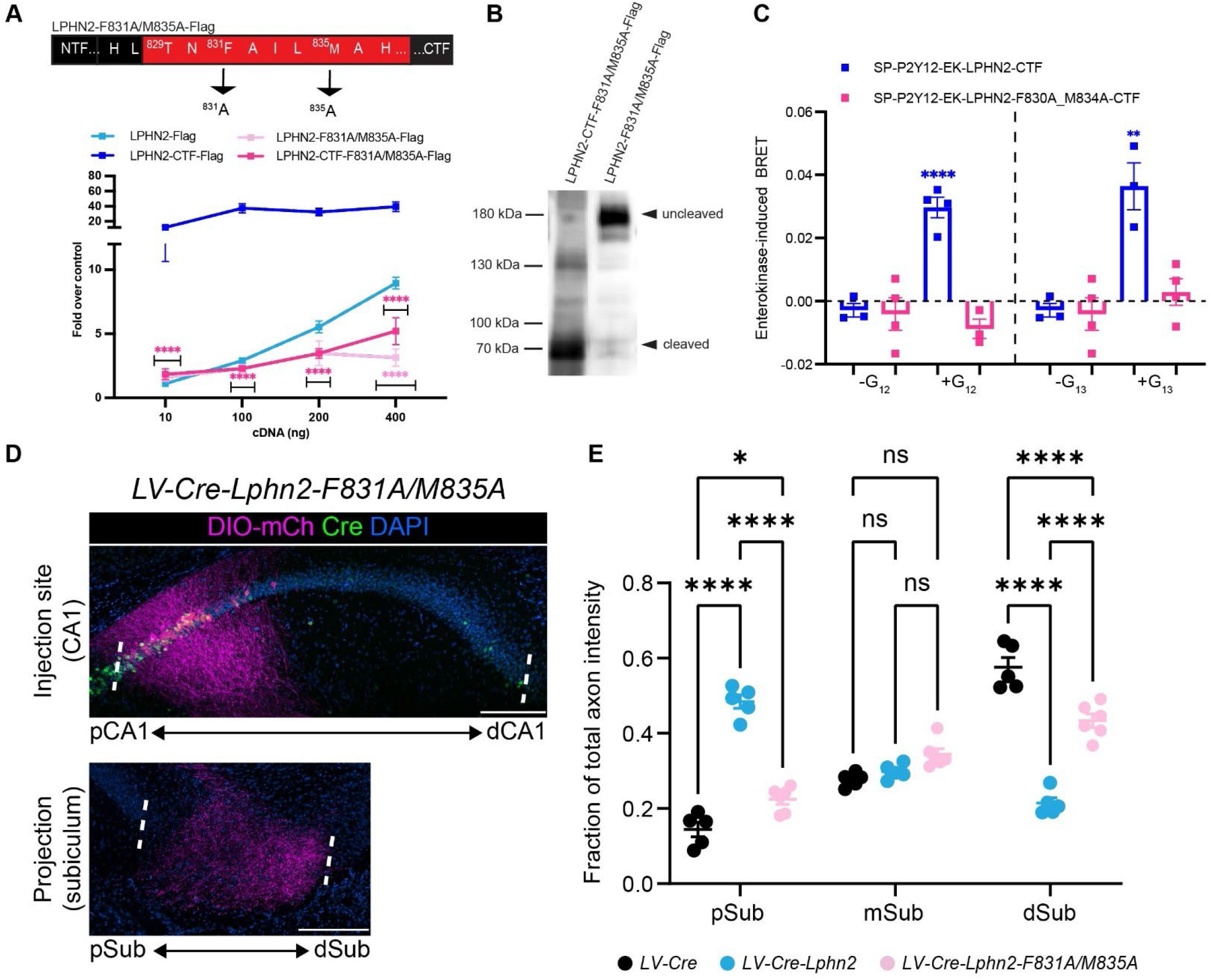
Lphn2-F831A/M835A has impaired G protein coupling activity and autoproteolytic cleavage and fails to misdirect pCA1 axons to the pSub when misexpressed. **(A)** Schematic of the mutated TA for Lphn2-F381A/M835A. The SRE luciferase reporter assay shows that both the FL Lphn2-F831A/M835A and the Lphn2-F831A/M835A-CTF have impaired signaling. Means ± SEM; Multiple unpaired t tests between FL Lphn2 and Lphn2-F831A/M835A and Lphn2-CTF and Lphn2-F831A/M835A-CTF constructs; ****, p<0.0001. **(B)** Immunoblot analysis of TA-dead Lphn2 and TA-dead Lphn2-CTF expression in HEK293T cells using a primary antibody against Flag (1:500, ThermoFisher, PA1-984B). Expected bands for FL Lphn2-F831A/M835A-Flag and Lphn2-F831A/M835A-CTF-Flag are 164 kDa and 72 kDa, respectively. **(C)** Gβγ-release BRET assay testing SP-P2Y12-EK-Lphn2-F831A/M835A-CTF activation of Gα_12_ and Gα_13_ in HEKΔ7 cells. SP-P2Y12-EK-Lphn2-CTF signaling is shown for comparison. Means ± SEM; Multiple unpaired t tests between no G protein and G protein conditions; **, p<0.01; ****, p<0.0001. **(D)** Representative images of *AAV-DIO-mCh* (magenta; mCh expression in a Cre-dependent manner) injections in proximal CA1 (top) and corresponding projections in the subiculum (bottom). **(E)** Fraction of total axon intensity within proximal, mid, and distal subiculum. *LV-Cre*: n=5, *LV-Cre-Lphn2*: n=5 and *LV-Cre-Lphn2-F831A/M835A*: n=6. Means ± SEM; two-way ANOVA with Sidak’s multiple comparisons test. Injection sites of all subjects are shown in **Figure 1—figure supplement 1**. Scale bars represent 200 μm.

To confirm that the reduced SRE response was due to impaired G protein coupling and not simply to impaired proteolysis, we cloned the CTF of our Lphn2-F831A/M835A mutant into our enterokinase-activatable construct. We then tested our construct in the Gβγ-release assay and compared the BRET response to WT Lphn2-CTF. Unlike the WT receptor, Lphn2-CTF-F831A/M835A did not yield a BRET signal after re-introduction of any of the G proteins in question (Gα_s_, Gα_i1,_ Gα_q_, Gα_12_, or Gα_13_) (**Figure 3C, Figure 2—figure supplement 2**). Taken together, our findings demonstrate that the F831A/M835A mutations in Lphn2 impair TA-mediated G protein coupling.

### TA activity or autoproteolysis of Lphn2 is required for its cell-autonomous effect in causing proximal CA1 axon mistargeting

We next misexpressed *Lphn2-F831A/M835A* in proximal CA1 to determine if Lphn2 TA activity or autoproteolysis is required *in vivo* to direct mistargeting of proximal CA1 axons. We injected *LV-Cre-Lphn2-F831A/M835A* into CA1 of P0 mice, followed by *AAV-DIO-mCherry* into proximal CA1 of the same mice as adults. The majority of Lphn2-F831A/M835A-expressing proximal CA1 axons targeted the most distal third of the subiculum, like negative control *LV-Cre* animals (**Figure 3D**). The fraction of axon intensity in *LV-Cre-Lphn2-F831A/M835A* animals was significantly lower in the proximal subiculum and significantly higher in distal subiculum when compared to *LV-Cre-Lphn2* animals (**Figure 3E**). Proximal CA1 axons in *LV-Cre-Lphn2-F831A/M835A* animals showed a similar pattern of targeting to negative control *LV-Cre* animals (**Figure 3—figure supplement 2**), although the total fraction of axon intensity was significantly lower in the distal subiculum (**Figure 3E**).

Collectively, these findings suggest that TA activity or autoproteolysis is required for Lphn2-mediated miswiring of proximal CA1 axons.

### Mutating residue T829G in the TA renders Lphn2 cleavage deficient but preserves the ability of the TA to activate G protein

While misexpressing Lphn2-F831A/M835A failed to cause proximal CA1 axons to mistarget to the proximal subiculum, we could not definitively link this result to impaired TA activity since the Lphn2-F831A/M835A mutant was also resistant to autoproteolytic cleavage (**Figure 3B**). Since our initial efforts to find a TA mutant with impaired G protein signaling that retained normal autoproteolytic cleavage were unsuccessful, we designed a construct that rendered Lphn2 resistant to autoproteolytic cleavage but preserved TA activity. Previous studies showed that replacing threonine-838 in the TA of Lphn1 or threonine-923 in the TA of Lphn3 to glycine inhibited autoproteolysis while maintaining proper folding of the receptor (Araç et al., 2012; Kordon et al., 2022). Thus, we mutated the analogous threonine-829 in Lphn2 (Lphn2-T829G) and confirmed that Lphn2-T829G was cleavage resistant using immunoblotting (**Figure 4A**). We also validated that Lphn2-T829G is expressed on the cell surface at a comparable level as Lphn2 WT (**Figure 3—figure supplement 1**).

**Figure 4.**
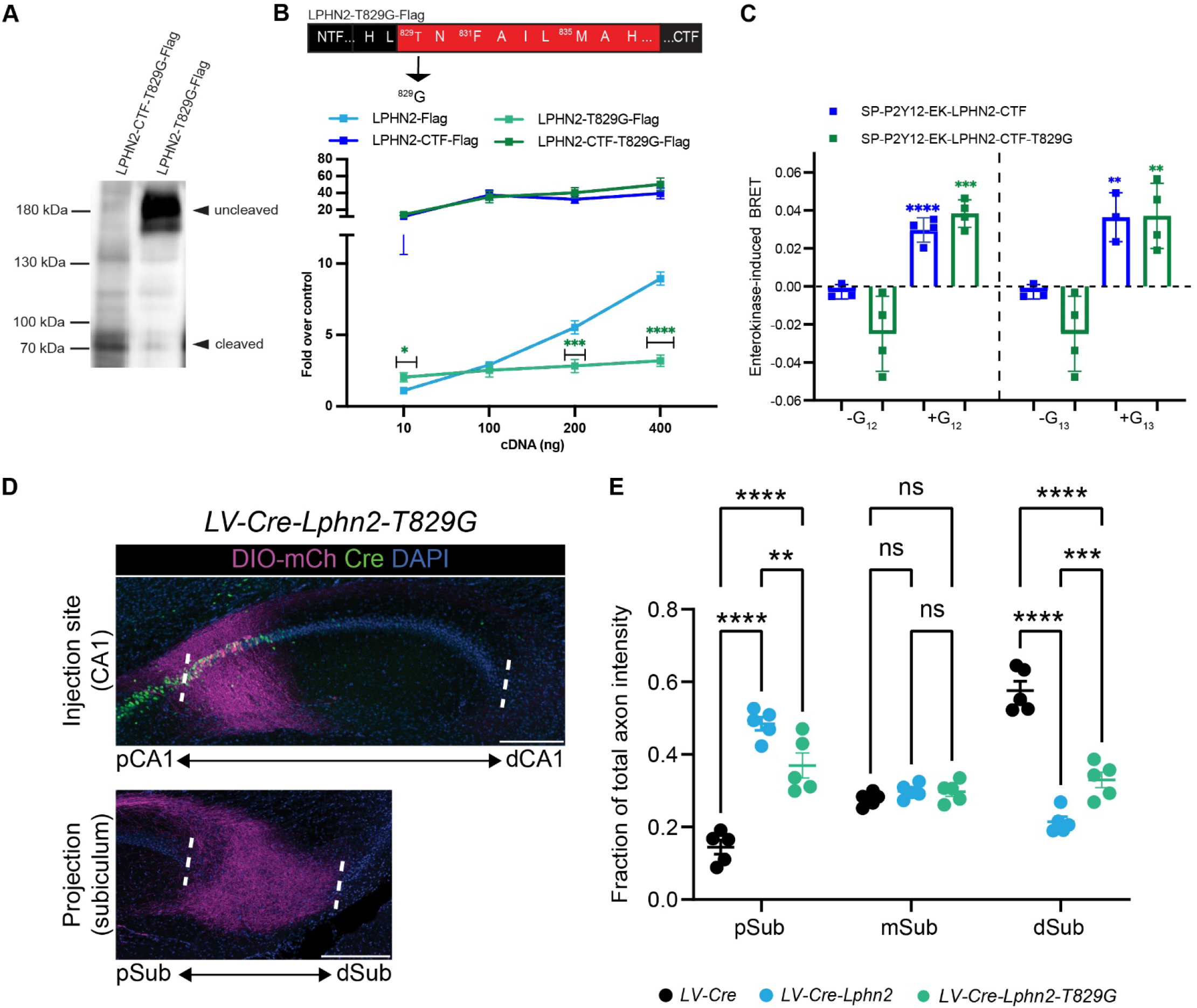
Lphn2-T829G impairs autoproteolytic cleavage, retains G protein activity in the truncated receptor, and misdirects axons to the pSub when misexpressed. **(A)** Immunoblot analysis of Lphn2-T829G and Lphn2-T829G-CTF expression in HEK293T cells using a primary antibody against Flag (1:500, ThermoFisher, PA1-984B). Expected bands for FL Lphn2-T829G-Flag and Lphn2-T829G-CTF-Flag are 164 kDa and 72 kDa, respectively. **(B)** Schematic of the mutated TA for Lphn2-T829G. The SRE luciferase reporter assay shows that the FL Lphn2-T829G has impaired SRE levels while the Lphn2-T829G truncated up to the GPS cleavage site has SRE levels comparable to Lphn2-CTF. Means ± SEM; Multiple unpaired t tests between FL Lphn2 and Lphn2-T829G and Lphn2-CTF and Lphn2-T829G-CTF constructs; *, p<0.05; ***, p<0.001; ****, p<0.0001. **(C)** Gβγ-release BRET assay testing SP-P2Y12-EK-Lphn2-T829G-CTF activation of Gα_12_ and Gα_13_ in HEKΔ7 cells. SP-P2Y12-EK-Lphn2-CTF signaling is shown for comparison. Means ± SEM; Multiple unpaired t tests between no G protein and G protein conditions; **, p<0.01; ***, p<0.001; ****, p<0.0001. **(D)** Representative images of *AAV-DIO-mCh* (magenta; mCh expression in a Cre-dependent manner) injections in proximal CA1 (top) and corresponding projections in the subiculum (bottom). **(E)** Fraction of total axon intensity within proximal, mid, and distal subiculum. *LV-Cre*: n=5, *LV-Cre-Lphn2*: n=5 and *LV-Cre-Lphn2-T829G*: n=5. Means ± SEM; two-way ANOVA with Sidak’s multiple comparisons test. Injection sites of all subjects are shown in **Figure 1—figure supplement 1**. Scale bars represent 200 μm.

We next assessed G protein signaling for Lphn2-T829G using our SRE gene expression system with FL and truncated receptors (Lphn2-T829G and Lphn2-CTF-T829G, respectively) (**Figure 4B**). FL Lphn2-T829G had significantly impaired SRE response compared to WT Lphn2, consistent with diminished exposure of the TA in the absence of cleavage; however, when we tested Lphn2-CTF-T829G, which lacked the entire NTF up to the GPS cleavage site, we observed SRE levels comparable to Lphn2-CTF suggesting that the mutated TA is fully active if exposed. We therefore cloned the CTF of the T829G mutant into our enterokinase-activatable construct and tested BRET signaling following re-introduction of Gα proteins (**Figure 4C**). The T829G-CTF retained BRET signaling comparable to WT Lphn2-CTF for Gα_12_ and Gα_13_, with no discernable BRET response for Gα_s_, Gα_i1,_ or Gα_q_ (**Figure 2—figure supplement 2**). Taken together, these data supported that Lphn2-T829G is resistant to autoproteolytic cleavage but maintains a functional TA.

### Autocleavage-deficient Lphn2 retains moderate activity in directing proximal CA1 axon mistargeting

To assess if autoproteolytic cleavage is required *in vivo* for Lphn2-mediated proximal CA1 axon mistargeting, we injected *LV-Cre-Lphn2-T829G* into proximal CA1 of P0 mice, followed by *AAV-DIO-mCherry* into pCA1 of the same mice as adults. Overall, Lphn2-T829G-expressing proximal CA1 axons did not show highly enriched targeting to a specific region of the subiculum, as observed for control proximal CA1 axons or Lphn2 misexpressing proximal CA1 axons, which preferentially target distal and proximal subiculum, respectively (**Figure 4E**). However, compared to control there was a significant increase in the fraction of axon intensity in proximal subiculum compared to mid subiculum and distal subiculum in *LV-Cre-Lphn2-T829G* animals, although this mistargeting was not as pronounced as seen with WT *LV-Cre-Lphn2* (**Figure 3—figure supplement 2**.).

The intermediate gain-of-function phenotypes of misexpressing Lphn2-T829G compared to misexpressing WT Lphn2 or TA-deficient (and non-cleavable) Lphn2 suggest that autoproteolysis is not absolutely required for Lphn2 misexpression-induced miswiring of proximal CA1 axons. The weaker phenotype than WT is likely caused by the decreased G protein signaling of the FL construct given the more limited exposure of the TA in the absence of cleavage. Thus, misexpressing *LV-Cre-Lphn2-T829G* compared to *LV-Cre-Lphn2-F831A/M835A* gave a targeting phenotype closer to that of WT Lphn2 misexpression, consistent with the ability of its TA to signal.

### Neither TA activity nor autoproteolysis is required for Lphn2’s action as a repulsive ligand

We previously showed that misexpression of Lphn2 in distal subiculum target neurons causes proximal CA1 axons to avoid this area, suggesting that Lphn2 acts cell non-autonomously as a repulsive ligand in directing target selection of proximal CA1 axons (Pederick et al., 2021). In the context of repulsive axon guidance, proteolysis has been proposed as a mechanism to disassemble the extracellular binding complex after repulsive signaling, which is necessary for repulsion (Hattori et al., 2000). Are TA activity and/or autoproteolysis required for Lphn2’s cell non-autonomous role in neural circuit assembly? To test this, we used a dual injection strategy to ectopically express Lphn2 in distal subiculum and trace proximal CA1 axons into the subiculum (**Figure 5—figure supplement 1)**. At postnatal day 0 (P0), lentivirus expressing *GFP* alone (*LV-GFP*), or *GFP* and Lphn2 variants (*LV-Lphn2*, *LV-Lphn2-F831A/M835A* and *LV-GFP-Lphn2-T829G*) was injected into distal subiculum, followed by injection of membrane-bound mCherry (*AAV-mCherry*) into proximal CA1 in the same mice at approximately P42 (**Figure 5A**). The P0 lentivirus injection only covers a small fraction of the entire proximal CA1 axon projection, enabling us to assess whether proximal CA1 axons target lentivirus-expressing regions differently to adjacent regions that do not express lentivirus. To observe the relationship between proximal CA1 axon projections and lentivirus induced regions of the subiculum, we plotted axon signal intensity (mCh) and lentivirus injection site (GFP) from the same animal as height and color, respectively.

**Figure 5.**
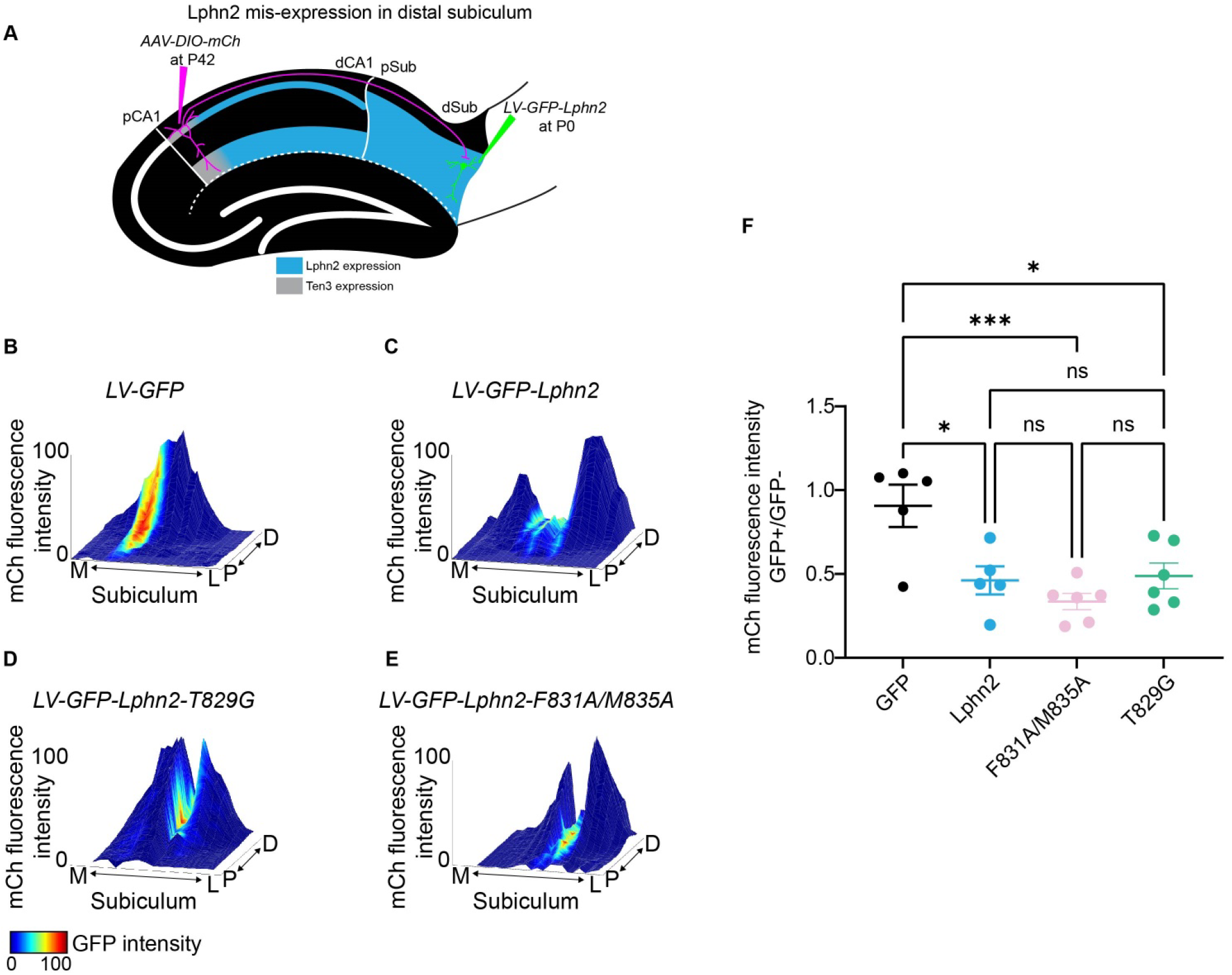
Misexpression of Lphn2 mutants in target distal subiculum neurons does not cause mistargeting of proximal CA1 axons. **(A)** Experimental design of Lphn2 misexpression assay in distal subiculum. **(B** to **E)**. Representative mountain plots showing normalized mCh fluorescence as height (proximal CA1 axon projections in subiculum) and normalized GFP fluorescence as color (lentivirus expression). P, proximal; D, distal; M, medial; L, lateral. **(F)** Ratio of mCh fluorescence intensity (from proximal CA1 axons) in GFP+ versus GFP– regions of the subiculum. *LV-GFP*: n=5, *LV-GF{-Lphn2*: n=5, *LV-Cre-Lphn2-F831A/M835A*: n=6 and *LV-GFP-Lphn2-T829G*: n=6. Means ± SEM. One-way analysis of variance (ANOVA) with Tukey’s multiple comparisons test. Data from (B), (C) and corresponding data in (F) are from a previous publication (Pederick et al., 2021).

As we previously described, GFP alone does not affect the intensity of proximal CA1 axons, whereas GFP-Lphn2 regions have significantly reduced proximal CA1 axon intensity in GFP-Lphn2 positive regions ((Pederick et al., 2021) and **Figures 5B, C, and F**). When either *LV-Lphn2-F831A/M835A* or *LV-GFP-Lphn2-T829G* were expressed in distal subiculum, we also observed a significant decrease in axon intensity in GFP positive regions compared to *LV-GFP* (**Figures 5D, E, and F**). This decrease was not significantly different from *LV-GFP-Lphn2* animals (**Figure 5F**). These findings suggest that neither TA activity nor autoproteolysis is required for Lphn2’s cell non-autonomous role as a ligand in neural circuit assembly.

## DISCUSSION

In this study, we utilized a combination of *in vivo* axon target selection and *in vitro* cell signaling assays to determine if Lphn2 G protein signaling is required for its role as a neural wiring molecule. First, we showed that Lphn2 misexpression can cell-autonomously misdirect proximal CA1 axons to proximal subiculum, establishing an assay to test the requirements of Lphn2 G protein signaling when it acts as a receptor (**Figure 1**). Second, we identified the G protein interaction partners of Lphn2 (**Figure 2**) and validated point mutations that disrupt TA activity and/or autoproteolysis of the GPS region (**Figures 3, 4**). Third, we showed that when Lphn2 is misexpressed in CA1 axons TA activity is required for Lphn2’s ability to misdirect axon targeting (**Figures 3, 4**). Finally, when Lphn2 acts as a repulsive ligand in subiculum target neurons, we demonstrated that neither TA activity nor autoproteolytic cleavage is required for the receptor’s ability to repel Ten3+ proximal CA1 axons (**Figure 5**). Taken together, these findings highlight the importance of Lphn2 G protein signaling during precise circuit assembly in a context-specific manner. Our results also support that while aGPCR GPS cleavage is dispensable Lphn2’s role as a receptor to direct axon targeting, an intact TA is essential.

### The role of autoproteolytic cleavage and tethered agonism in aGPCR activation

Upon aGPCR biosynthesis, the conserved GAIN domain undergoes autoproteolytic cleavage at the GPS to generate N- and C-terminal fragments that remain non-covalently bound during trafficking to the cell surface (Araç et al., 2012). Crystal structures of GAIN domains from Lphn1 and ADGRB3 (BAI3) (Araç et al., 2012), ADGRG1 (GPR56) (Salzman et al., 2016), and ADGRG6 (GPR126) (Leon et al., 2020), revealed that in an intact aGPCR the TA is buried as a ꞵ-strand in the GAIN domain, forming an extensive network of conserved hydrogen bonds and hydrophobic side chains. This suggests that when the complex between aGPCR’s NTF and CTF remains intact the TA is inaccessible for 7TM domain engagement. However, several studies have reported that naturally cleavage-resistant aGPCRs can still function (Liebscher et al., 2014; Wilde et al., 2016). This suggests that the TA can in fact interact with the 7TM independent of cleavage, although the efficacy of this interaction is likely to be diminished, as we see in the present study. Thus, the relative contributions and/or necessity of autoproteolytic cleavage and the TA to aGPCR activity remain an area of active study.

Recent efforts reported the structures of 8 aGPCRs, 7 of which were truncated up to the GPS (Barros-Álvarez et al., 2022; Ping et al., 2022; Qu et al., 2022; Xiao et al., 2022). While most discussions from the structural aGPCR studies argued that NTF dissociation is required for TA interaction with the receptor, the structure of autoproteolysis-deficient ADGRF1 supported a cleavage-independent manner of receptor activation (Qu et al., 2022). The density for the TA of ADGRF1 was well-resolved and bound in an ɑ-helical structure within the orthosteric site of the 7TM bundle. This interaction was like that of the cleaved structures, and ADGRF1 was also observed to be bound to a miniG_i1_, supporting that receptor cleavage and TA exposure are not absolutely required for G protein coupling.

As mentioned above, not all aGPCRs are auto-proteolytically cleaved; therefore, activation cannot be fully dependent on TA exposure through removal of the NTF (Kishore et al., 2016; Liebscher et al., 2022). In this regard, it is possible that FL aGPCRs exist in multiple conformational states that include receptor molecules in which the TA is unmasked from the GAIN domain. In fact, molecular dynamics (MD) simulations of spontaneous TA exposure were recently reported for five intact aGPCR homologs (ADGRB3, ADGRE2, ADGRE5, ADGRG1, and Lphn1) (Beliu et al., 2021). Here, the authors show that TA exposure occurs due to the high intrinsic flexibility of the GAIN domain. They also used biorthogonal labeling of conserved positions within the TA to show that large portions (+6 residues) of the TA can become solvent accessible in the context of the GAIN domain. They argue that TA exposure likely occurs in a stepwise mechanism where the TA is uncovered along its N→C axis. Thus, it is possible that an intact complex of aGPCR’s NTF and CTF could unmask the TA sufficiently for interaction with the 7TM, resulting in receptor activation.

The ability of TA exposure to occur in intact aGPCRs could provide an explanation for why our Lphn2-T829G mutant displays a partial axon mistargeting phenotype (**Figure 4**). Even though the Lphn2-T829G mutant cannot undergo autoproteolytic cleavage, it still retains a functional TA that is able to initiate G protein signaling if transiently unmasked. This is likely why activation of FL Lphn2-T829G is less robust than the WT Lphn2, which can more readily unmask the TA.

Like the Lphn2-T829G mutant, Lphn2-F831A/M835A cannot undergo autoproteolytic cleavage. However, this mutant also has impaired G protein coupling activity even with full exposure of the TA. This explains why we observed essentially normal axon targeting to the distal subiculum when we overexpressed Lphn2-F831A/M835A in proximal CA1 (**Figure 3**). Even if the TA of Lphn2-F831A/M835A becomes unmasked it still cannot initiate TA-mediated receptor activity.

### Implication of Lphn2 signaling in neural circuit assembly

How could Lphn2-mediated G protein signaling lead to axon repulsion? We show here that Lphn2 primarily signals through Gα_12_ and Gα_13_ (**Figure 2E**). Gα_12_/Gα_13_ are known to regulate Rho GTPase; for example, Gα_13_ binds to and activates p115RhoGEF, an exchange factor for and activator of the small GTPase RhoA (Kozasa et al., 2011). RhoA activation is known to cause growth cone collapse and neuronal process retraction via its regulation of the actin-myosin contractility (Luo, 2002; Spillane and Gallo, 2014). Thus, Gα_12_/Gα_13_→RhoGEF→RhoA may be a plausible pathway for Lphn2 to mediate its function as a receptor for axon repulsion.

Interestingly, neither autoproteolytic cleavage nor TA activity is required for Lphn2 to act cell non-autonomously as a repulsive ligand in target neurons (**Figure 5**). This suggests that cleavage of Lphn2 is not required for repulsion in this context and implies that another mechanism mediates disassembly of the extracellular binding complex, which is required for retracting axons to pull away from the targets. Other potential mechanisms to disassemble the extracellular binding complex include Ten3 cleavage (teneurins are known to also undergo proteolytic cleavage at its extracellular domain; Sita et al., 2019), endocytosis of the adhesion complex as in the case of ephrin/Eph receptor (Egea and Klein, 2007), or forces produced by actin-myosin contractility in axon terminals induced by repulsion signaling. Indeed, extracellular binding of Lphn2 to Ten3 should in principle trigger a repulsive response in Ten3-expressing axon terminals, but the signaling mechanism is completely unknown. Future studies on the mechanisms that disassemble the extracellular complex and intracellular signaling downstream of Ten3 will increase our understanding of how interaction of two molecules can lead to repulsive outcomes.

## MATERIALS AND METHODS

### Materials for cell culture experiments

Dulbecco’s Modified Eagle Medium (DMEM), high glucose, and penicillin-streptomycin (P/S) (10,000 U/mL) were purchased from Gibco (ThermoFisher Scientific, Waltham, MA). Fetal bovine serum (FBS), 0.5% trypsin, and Dulbecco’s phosphate-buffered saline (DPBS) were purchased from Corning (Fisher Scientific, Waltham, MA). Opti-MEM reduced serum medium, no phenol red, and Lipofectamine 2000 transfection reagent were purchased from Invitrogen (ThermoFisher Scientific). FuGENE transfection reagent was purchase from Promega (Madison, WI). RIPA buffer was purchased from Sigma-Aldrich (St. Louis, MO). Triton lysis buffer consisted of 0.11 M Tris-HCl powder, 0.04 M Tris-base powder, 75 mM NaCl, 3 mM MgCl_2_, and 0.25% Triton X-100 pure liquid. The 3X Firefly Assay Buffer was freshly prepared in Triton lysis buffer and contained 15 mM DTT, 0.6 mM coenzyme A (MedChemExpress, Monmouth Junction, NJ), 0.45 mM ATP (MedChemExpress, Monmouth Junction, NJ), and 0.42 mg/mL firefly D-luciferin (NanoLight Technology). *Renilla* Salts buffer consisted of 45 mM Na_2_EDTA, 30 mM Na Pyrophosphate, and 1.425 M NaCl. The 3X *Renilla* Assay Buffer was freshly prepared in *Renilla* Salts and contained 0.06 mM PTC124 in DMSO (MedChemExpress) and 0.01 mM coelenterazine-*h* (NanoLight Technologies, Pinetop, AZ). For the BRET assays, enterokinase, light chain, was obtained from New England Biolabs (Ipswich, MA), isoproterenol and quinpirole from Sigma Aldrich, and endothelin 1 (ET-1) from Tocris Bioscience (Bristol, United Kingdom). YM-254890 was purchased from AdipoGen Life Sciences (San Diego, CA). Impermeant Janelia Fluor 646 conjugated to benzyl guanine was a kind gift from Dr. Luke Lavis (Howard Hughes Medical Institute Janelia Research Campus).

### Plasmid DNA constructs

FL *Lphn2* was amplified from cDNA isolated from the P8 mouse hippocampus. This cDNA was used as a polymerase chain reaction (PCR) template to make the *Lphn2* constructs used in this study. All cDNA constructs were assembled in a pCDNA3.1+ vector by Gibson assembly using NEBuilder HiFi DNA Assembly Master Mix (New England Biolabs). Sequences were confirmed with Genewiz sequencing service (South Plainfield, NJ).

### Cell culture

HEK293T cells (American Type Culture Collection, Manassas, VA) and HEK293 cells with targeted deletion via CRISPR-Cas9 of *GNAS, GNAL, GNAQ, GNA11, GNA12, GNA13, and GNAZ* (HEKΔ7)(Alvarez-Curto et al., 2016) were maintained in high-glucose DMEM supplemented with 10% FBS and 1% P/S at 37°C in a 5% CO_2_ humidified incubator.

### Immunoblot analysis

HEK293T cells were detached for 2–3 min using 0.5% trypsin and then plated at a density of 350,000 cells/mL in a 6-well culture plate. After 24 hr, the cells were transfected using FuGENE transfection reagent (8 μL/2 μg cDNA) and Opti-Mem with receptor cDNA (2 μg). After 24 hr, cells were placed on ice and incubated in 500 μL RIPA buffer for 30 min. Following this incubation, cells were scraped from the culture plate and moved into 1.5 mL microcentrifuge tubes. Cells were then spun at 15,000 xg in a 4°C benchtop centrifuge to pellet debris. After centrifugation, 50 μL of the supernatant was transferred into a fresh microcentrifuge tube and combined with 50 μL 2X SDS Laemmli sample buffer (Sigma-Aldrich). In preparation for immunoblot analysis, 20 μL sample was run on an SDS-PAGE gel (Mini-PROTEAN TGX, 4-15%, Bio-Rad Laboratories, Inc., Hercules, CA) prior to transfer to a PDVF membrane (Immobilon-P Membrane, Merck Millipore Ltd., Burlington, MA). The membrane was then incubated in a 5% milk tris-buffered saline with 0.1% tween-20 (TBS-T) solution for 1 hr at RT with gentle rotation. The membrane was washed 5 times 5 min in TBS-T prior to overnight incubation at 4°C with 1° anti-Flag antibody (1:500, ThermoFisher, PA1-984B). The next morning, the membrane was washed 5 times 5 min in TBS-T. The membrane was then incubated for 1 hr at RT with 2° anti-rabbit HRP antibody (1:10,000, ThermoFisher, Cat #31458). The membrane was washed 5 times 5 min in TBS-T prior to visualization with SuperSignal West Pico Chemiluminescent Substrate (Fisher Scientific) using the Azure c600 Gel Imaging System (Azure Biosystems, Dublin, CA).

### Gene expression assays

In preparation for transfection, HEK293T cells were detached for 2–3 min using 0.5% trypsin and then seeded at a density of 400,000 cells/mL in a 12-well culture plate. After 24 hr, the cells were co-transfected using Lipofectamine 2000 (2.5 μL/1 μg cDNA) and Opti-Mem with receptor cDNA (10-600 ng), gene reporter cDNA (600 ng), and empty vector pCDNA5/FRT to balance the total amount of cDNA up to 1,200 ng. After 6 hr with the transfection reagent, the media was volume exchanged to serum-free DMEM supplemented with 1% P/S (∼18 hr serum starvation).

After 24 hr, the media was aspirated from the cells and each well was gently rinsed with DPBS. Cells were then mechanically detached using 275 μL DPBS and 80 μL of the resuspension was distributed in triplicate to a 96-well black/white isoplate (Perkin Elmer Life Sciences). Next, 40 μL of 3X Firefly Assay Buffer was added to each well. The emission was then read at 535 nm after 10 min incubation using a PHERAstar FS microplate reader (BMG LABTECH, Ortenberg, Germany). Next, 60 μL 3X *Renilla* Assay Buffer was added to each well. The emission was then read at 475 nm after 10 min incubation using a PHERAstar FS microplate reader. For assays using the Gαq-inhibitor YM-254890, the cell media was exchanged to DMEM containing 1 μM YM-254890 approximately 6 hr after transfection.

### Bioluminescence resonance energy transfer assays

In preparation for transfection, HEKΔ7 cells were detached for 2–3 min using 0.5% trypsin and then seeded at a density of 400,000 cells/mL in a 12-well culture plate. After 24 hr, the cells were co-transfected using Lipofectamine 2000 (2.5 μL/1 μg cDNA) and Opti-Mem with receptor cDNA (200ng), Gα (720 ng), Gβ1 (250 ng), Gγ2-Venus (250 ng), membrane-anchored GRK3ct-Rluc8 (50 ng), and empty vector pCDNA5/FRT to balance the total amount of cDNA up to 1,470 ng. After 24 hr transfection, cells were washed with DPBS before being re-suspended in 400 μL BRET buffer (DPBS containing 5 mM glucose). Next, 45 μL of the resuspension was distributed to six wells of a 96-well OptiPlate black-white plate (Perkin Elmer Life Sciences, Waltham, MA). Cells were then incubated for 10 mins with 10 μL coelenterazine-*h* (final concentration 5 μM) before ligand addition to reach a final well volume of 100 μL. Donor (Rluc8) and acceptor (mVenus) emission was read using a PHERAstar FS microplate reader at 485 nm and 525 nm, respectively. The BRET ratio was then measured by dividing the 525 emissions by the 485 emissions. The drug-induced BRET ratio was then calculated by subtracting the buffer BRET for each condition.

### Surface expression measurements using SNAPfast-tag

In preparation for transfection, HEK293T cells were detached for 2–3 min using 0.5% trypsin and then seeded at a density of 900,000 cells/well in a 6-well culture plate. After 24 hr, the cells were transfected using FuGENE transfection reagent (8 μL/2 μg cDNA) and Opti-Mem with SNAPfast-tagged receptor cDNA (2 μg). After 24 hr, cells were incubated for 30 min with 500 µL 1 µM impermeant Janelia Fluor 646 conjugated to benzyl guaanine was dissolved in DMEM containing 10% FBS and 1% P/S. Cells were then washed 3 times with complete DMEM and once with DPBS prior to resuspension in 500 µL DPBS. Next, 100 μL of resuspension was added to three wells of a 96-well OptiPlate black plate (Perkin Elmer Life Sciences, Waltham, MA). The emission was then read using the filter 640/680 at a gain of 1000 using a PHERAstar FS microplate reader (BMG LABTECH, Ortenberg, Germany).

### Mice

All procedures followed animal care and biosafety guidelines approved by Stanford University’s Administrative Panel on Laboratory Animal Care and Administrative Panel on Biosafety in accordance with NIH guidelines. Both male and female mice were used, and mice were group housed on a 12hr light/dark cycle with access to food and water *ad libitum*. CD-1 mice from Charles River Laboratories were used for all experiments. The total number of mice injected and screened for each experiment is as follows: Figure 1: *LV-Cre*, 101, *LV-Cre-Lphn2*, 60; Figure 3: *LV-Cre-Lphn2-F831A/M835A*, 86; Figure 4: *LV-Cre-Lphn2-T829G*, 102; and Figure 5: *LV-GFP-Lphn2-F831A/M835A*, 79 and *LV-GFP-Lphn2-T829G*, 56.

### Lentivirus generation

All lentivirus constructs expressing Cre, GFP, Lphn2, Lphn2-F831A/M835A or Lphn2-T829G were made by inserting corresponding cDNA into the *LV-UbC* plasmid (Pederick et al., 2021) with a P2A sequence between the two ORFs. Cre and GFP was amplified from *LV-UbC-GFP-Cre* and FL *Lphn2* cDNA was isolated from a cDNA library made form mRNAs from P8 mouse hippocampus. *GFP* and *Lphn2* were inserted into *LV-UbC* with Gibson assembly cloning kit (NEB E5510S). The Lphn2-F831A/M835A and Lphn2-T829G mutations were made using Q5 mutagenesis (NEB, E0552S). All plasmids were sequenced verified before virus was produced. All custom lentiviruses were generated by transfecting 36 10-cm plates (HEK293T) with 4 plasmids (4.1R, RTR2, VSVg, and transfer vector containing gene of interests). Medium was collected 48 hrs later and centrifuge at 8400 relative centrifugal force (rcf) for 18 hrs at 4°C. Viral pellets were dissolved with PBS and further purified with a 20% sucrose gradient centrifugation at 80 000 rcf for 2 hours.

### Stereotactic injections in neonatal mice

P0 mice were anesthetized using hypothermia. CA1 injections were 1.0 mm lateral, 0.85 mm anterior and 0.8 mm ventral from lambda and subiculum injections were 1.3 mm lateral, 0.45 mm anterior and 0.8 mm ventral from lambda. 100 nl of lentivirus was injected at 100 nl/min at the following titers: *LV-UbC-Cre* (7 x 10^12^ copies per ml), *LV-UbC-Cre-P2A-Lphn2-FLAG* (2.4 x 10^12^ copies per ml), *LV-UbC-Cre-P2A-Lphn2-F831A/M831A-FLAG* (2.24 x 10^12^ copies per ml) and *LV-UbC-Cre-P2A-Lphn2-T828G-FLAG* (1.2 x 10^13^ copies per ml), *LV-UbC-GFP* (6 × 10^12^ copies per ml), *LV-UbC-GFP-P2A-Lphn2-FLAG* (5 × 10^12^ copies per ml), *LV-UbC-GFP-P2A-Lphn2-F831A/M831A-FLAG* (3.6 x 10^12^ copies per ml), *LV-UbC-GFP-P2A-Lphn2-T828G-FLAG* (9 x 10^12^ copies per ml), *LV-UbC-Cre* (7 x 10^12^ copies per ml), *LV-UbC-Cre-P2A-Lphn2-FLAG* (2.4 x 10^12^ copies per ml), *LV-UbC-Cre-P2A-Lphn2-F831A/M831A-FLAG* (2.24 x 10^12^ copies per ml) and *LV-UbC-Cre-P2A-Lphn2-T828G-FLAG* (1.2 x 10^13^ copies per ml).

### Stereotactic injection in adult mice

Injections of *AAV8-EF1a-DIO-ChR2-mCh* (2 × 10^12^ copies per ml, Neuroscience Gene Vector and Virus core, Stanford University) and *AAV8-EF1a-ChR2-mCh* (2 × 10^12^ copies per ml, Neuroscience Gene Vector and Virus core, Stanford University) were performed at about P42. Mice were anesthetized using isoflurane and mounted in stereotactic apparatus (Kopf). Coordinates for proximal CA1 were 1.4 mm lateral and 1.25 mm posterior from bregma, and 1.12 mm ventral from brain surface. Virus was iontophoretically injected with current parameters 5 µA, 7 s on, 7 s off, for 2 min, using pipette tips with an outside perimeter of 10–15 μm. Mice were perfused about 2 weeks later and processed for immunostaining as described below.

### Immunostaining

Mice were injected with 2.5% Avertin and were transcardially perfused with PBS followed by 4% paraformaldehyde (PFA). Brains were dissected and post-fixed in 4% PFA overnight, cryoprotected for about 24 hours in 30% sucrose. Brains were embedded in Optimum Cutting Temperature (OCT, Tissue-Tek), frozen in dry ice cooled isopentane bath and stored at –80°C until sectioned. 60-μm thick floating sections were collected in PBS + 0.02% sodium azide and stored at 4°C. Sections were incubated in the following solutions at room temperature unless indicated: 1 hour in 0.3% PBS/Triton X-100 and 10% normal donkey serum, two nights in primary antibody at 4°C in 0.3% PBS/Triton X-100 and 10% normal donkey serum, 3 × 15 min in 0.3% PBS/Triton X-100, overnight in secondary antibody + DAPI (1:10,000 of 5 mg/ml, Sigma-Aldrich) in 0.3% PBS/Triton X-100 and 10% normal donkey serum, 2 × 15 min in 0.3% PBS/Triton X-100, and 15 min in PBS. Sections were mounted with Fluoromount-G (SouthernBiotech). Primary antibodies used were rat anti-mCherry (1:1,000, ThermoFisher, M11217, RRID:AB_2536611), rabbit anti-Cre (1:500, Synaptic Systems, 257 003, RRID:AB_2619968) and chicken anti-GFP (1:2,500, Aves Labs, GFP-1020, RRID:AB_10000240). Secondary antibodies conjugated to Alexa 488, Alexa 568, or Cy3 (Jackson ImmunoResearch) were used at 1:500 from 50% glycerol stocks.

### Image and data analysis for CA1 axon tracing

Mice were only included if they passed the following criteria: (1) AAV injection site must be in proximal CA1 (most proximal 30%), (2) lentivirus injections sites must be in CA1 and not in subiculum (Figure 1, 3 and 4) or lentivirus injections must be in the distal subiculum (Figure 5), (3) proximal CA1 axons must overlap with lentivirus injection site in subiculum (Figure 5). All mice that fulfilled these criteria are reported in Fig. 1, 3, 4 and 5 and were included in quantifications. Images of injections sites (5× magnification) and projections (10× magnification) were acquired of every other 60-μm sagittal section using a Zeiss epifluorescence scope. Due to variation in injection sites within each mouse, exposure was adjusted for each mouse to avoid saturation. Fluorescence intensity measurements on unprocessed images were taken using FIJI and data processing was performed using MATLAB.

For injection site quantification, a 30-pixel-wide segmented line was drawn from proximal CA1 to distal CA1 using DAPI signal as a guide. For projection quantification in subiculum, a 200-pixel-wide segmented line was drawn from proximal subiculum to distal subiculum through the cell body layer using only DAPI as a guide. From this point, injection site and projection images were processed the same. Segmented lines were straightened using the “Straighten” function, background subtraction was performed using the “Subtract” function and intensity values were measured using the “Plot Profile” command (FIJI). For injections that labeled both CA2 and proximal CA1, CA2 axons were present near the distal border of CA1 and spilled into proximal subiculum. These axons had their intensity set to zero by using area selection and the clear function (FIJI). The intensity plots were resampled into 100 equal bins using a custom MATLAB code.

For trace quantification in Figures 1, 3 and 4 the axon intensity was combined for all sections by summing all intensity values at each binned position. To calculate the fraction of axon intensity across proximal, mid, and distal subiculum the total axon intensity in bins 1–33, 34–66, and 67–100 was summed, respectively. The summed value from each of these regions was then divided by the total sum of axons from bins 1–100 to obtain the fraction of axons intensity within proximal, mid, and distal subiculum. Fractions of axon intensities were compared using a two-way ANOVA with Sidak’s multiple comparisons test using Prism 9 (GraphPad). The mean position of the injection sites was calculated by generating a summed intensity trace as above and then multiplying the intensity value by the bin position, summing across the entire axis, and dividing by the sum of the intensity values. Representative images (Figures 1, 3 and 4) were taken using a Zeiss LSM 780 confocal microscope (20× magnification, tile scan, max projection).

In Figure 5, the experimental and data analysis procedures were identical to Pederick et al., 2021 and therefore we used *LV-UbC-GFP* and *LV-UbC-GFP-P2A-Lphn2-FLAG* from that study to compare with *LV-UbC-GFP-P2A-Lphn2-F831A/M831A-FLAG* and *LV-UbC-GFP-P2A-Lphn2-T828G-FLAG* data generated from this study.

To quantify average axon intensity in GFP+ and adjacent GFP– regions in subiculum targets (Figure 5F), we restricted analysis to the most distal 20% of the subiculum. To determine the GFP+ region we identified the intensity-weighted central row using the summed fluorescence of each row and determined the minimal symmetric window of rows around the central row that encompassed at least 50% of the total intensity in the restricted GFP image. This defined a rectangle in the original image that we designated as the GFP+ region. We then computed the mean fluorescence intensity in this region for the mCh channel. We used the two rows above and below (lateral and medial) the designated GFP+ region as the adjacent GFP– region and computed the mean mCh fluorescence across these four rows. To determine mCh fluorescence differences in GFP+ versus GFP– regions, we divided the mCh intensity in the GFP+ region by the mCh intensity in the GFP– region for each mouse (i.e., GFP+/GFP–). mCh fluorescence intensity GFP+/GFP– was compared across groups using a one-way ANOVA with Tukey’s multiple comparisons test using Prism 9 (GraphPad). Three-dimensional mountain plots were generated using the ‘surf’ function.

## ACKNOWLEDGMENTS

We thank Z. Li, T. Li, C. McLaughlin, D. Wang, and Y. Wu for critiques of the manuscript. This work was support by NIH grants T32-MH015144 (NAPH), R01-NS050835 (LL), R01-MH54137 (JAJ), the Hope for Depression Research Foundation (JAJ) and P30EY012196 (ZH). L.L. is an investigator of the Howard Hughes Medical Institute.

## AUTHOR CONTRIBUTIONS

DTP, NAPH, JAJ, and LL conceived the study and designed the analysis. Data collection and analysis was performed by DTP and NAPH. HM and ZH generated custom lentivirus. DTP and NAPH wrote the manuscript with input from JAJ and LL. All authors approved the final version of the manuscript.

## MATERIAL AVAILABLILITY

All materials are available through requests to the corresponding author. All custom code was identical to reported in (Pederick et al., 2021) and can be accessed at https://github.com/dpederick/Reciprocal-repulsions-instruct-the-precise-assembly-of-parallel-hippocampal-networks/tree/1. All data generated or analyzed during this study are included in the manuscript and supporting file; Source Data files have been provided for all Figures.

## COMPETING INTERESTS

The authors declare that there are no relevant financial or non-financial competing interests to report.

## SUPPLEMENTARY FIGURES

**Figure 1—figure supplement 1.**
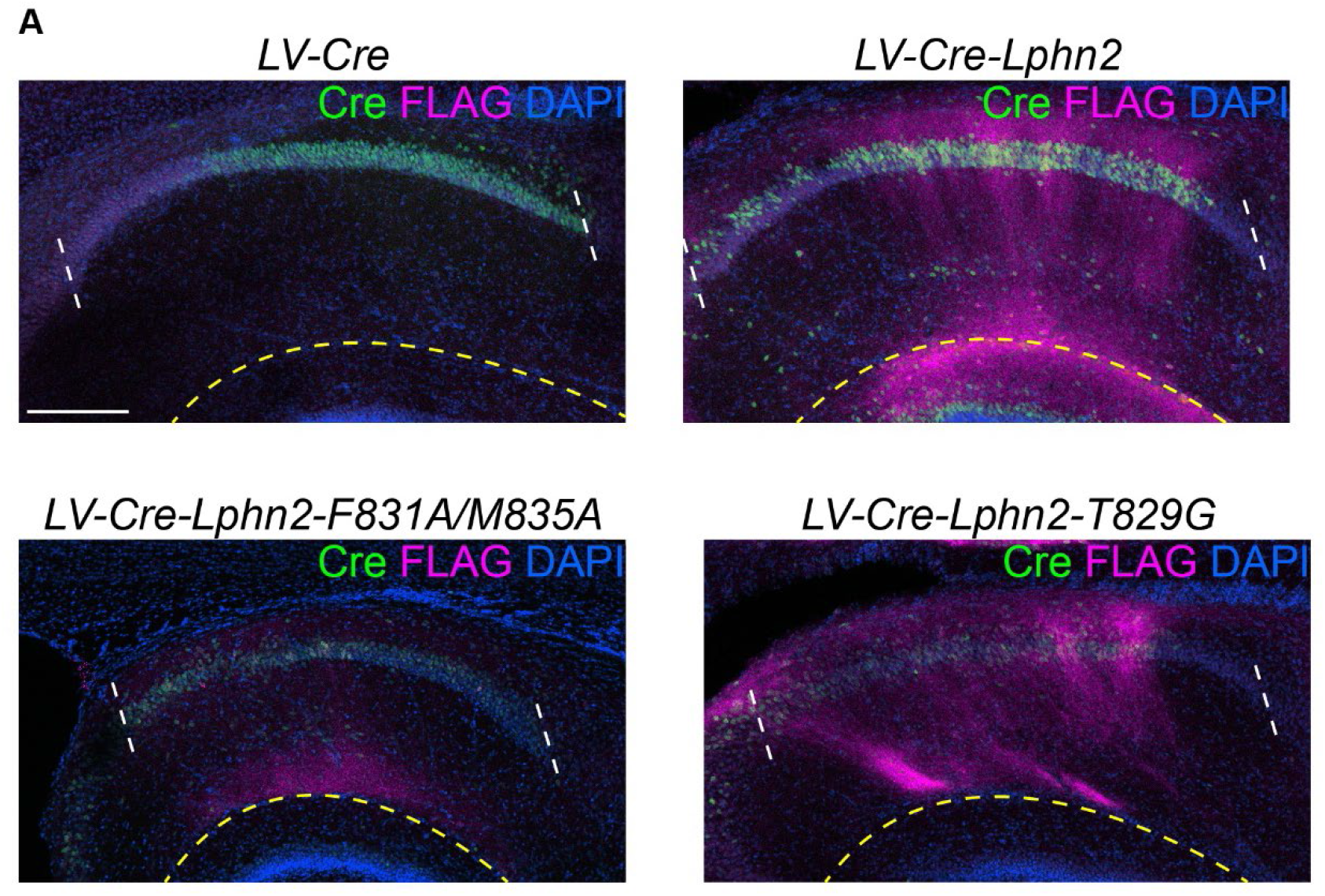
*In vivo* expression of lentivirus used in Figures 1, 3 and 4. (**A**) Representative images of Cre and FLAG immunostaining in P9 CA1 of mice injected with lentiviruses used in Figures 1, 3 and 4. The region between the white dashed line is CA1. The region below the yellow dashed line is the molecular layer of dentate gyrus. FLAG staining below the dashed yellow line is coming from dentate gyrus and not CA1 infected cells. Scale bar represents 200 μm.

**Figure 1—figure supplement 2.**
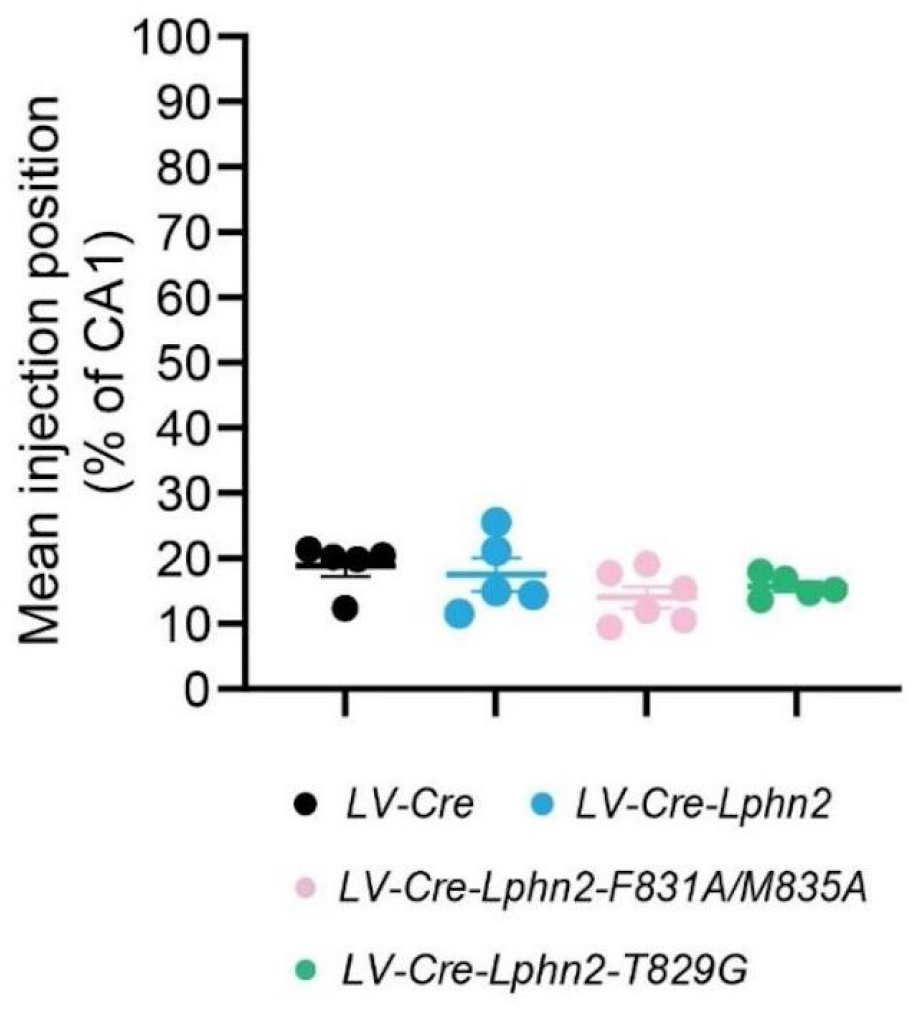
Mean injection site positions for proximal CA1 axon tracing in Figures 1, 3, and 4. All injection sites are in the most proximal 30% of CA1. The most proximal 30% of CA1 is the Lphn2-low region of CA1 (Pederick et al., 2021) and was therefore targeted for Lphn2 misexpression.

**Figure 2—figure supplement 1.**
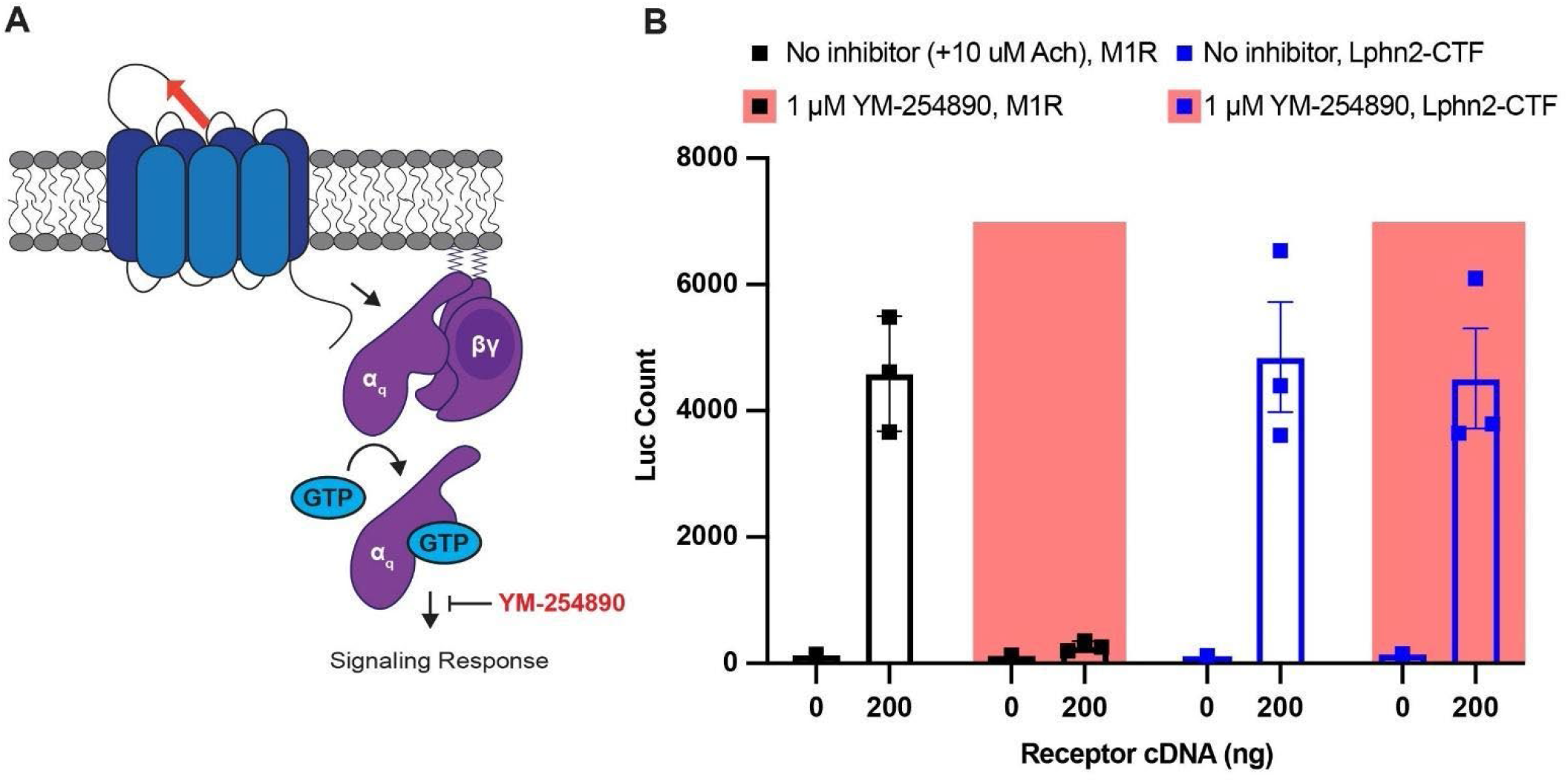
Gα_q_-inhibitor YM-254890 does not impair SRE luciferase response of Lphn2-CTF. **(A)** Schematic of YM-254890 activity in the gene reporter assay. YM-254890 inhibits signaling responses downstream of Gα_q_ activation. **(B)** 1 µM YM-254890 inhibited muscarinic acetylcholine receptor M1 (M1R) signaling in the SRE luciferase assay. For the M1R, 10 µM acetylcholine (ACh) was added for 6 hrs prior to reading the luminescence signal. 1 µM YM-254890 did not inhibit Lphn2-CTF signaling in the SRE luciferase assay.

**Figure 2—figure supplement 2.**
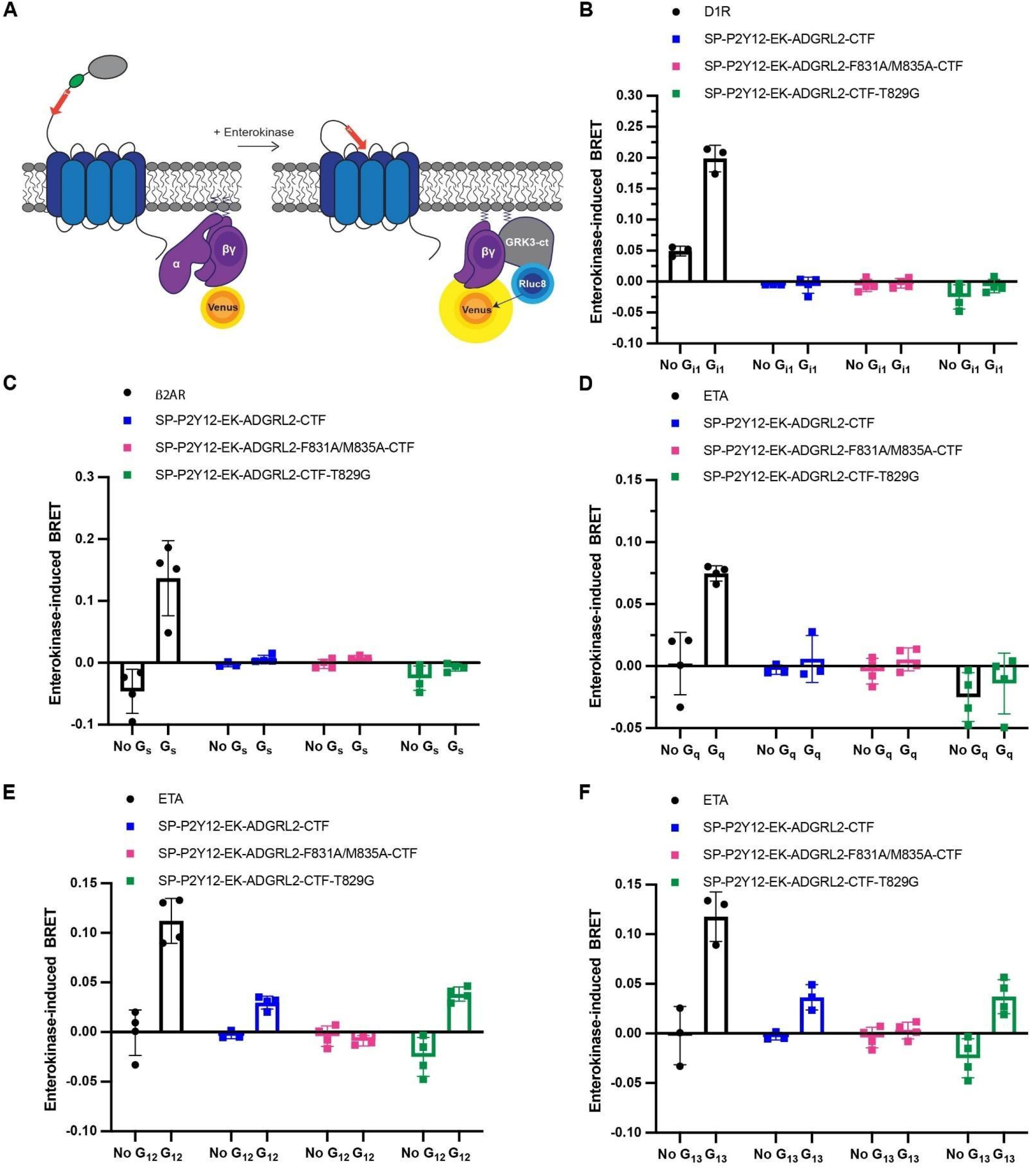
Gβγ-release BRET assay shows Lphn2 couples to Gα_12_ and Gα_13_. **(A)** Schematic outlining the Gβγ-release BRET assay. The Lphn2 TA is capped with an enterokinase cleavage site (EK) preceded by a hemagglutinin signal peptide (SP), P2Y12 N-terminal extracellular sequence, and flexible linker (Lizano et al., 2021). Addition of 10 nM enterokinase generates a TA neoepitope identical to activated endogenous Lphn2. Lphn2 activation results in G protein dissociation, allowing Gβγ-Venus to associate with the C-terminus of GPCR kinase 3 (GRK3-ct) (Hollins et al., 2009). Testing activation of SP-P2Y12-EK-Lphn2-CTF constructs with **(B)** Gα_i,_ **(C)** Gα_s_, **(D)** Gα_q_, **(E)** Gα_12_, and **(F)** Gα_13_ in HEKΔ7 cells. Dopamine receptor D2 (D2R) with 10 µM quinpirole, ꞵ2-adrenergic receptor (β2AR) with 1 µM isopropanol, and endothelin receptor (ETA) with 100 nM ligand ET-1 were used as positive controls.

**Figure 3—figure supplement 1.**
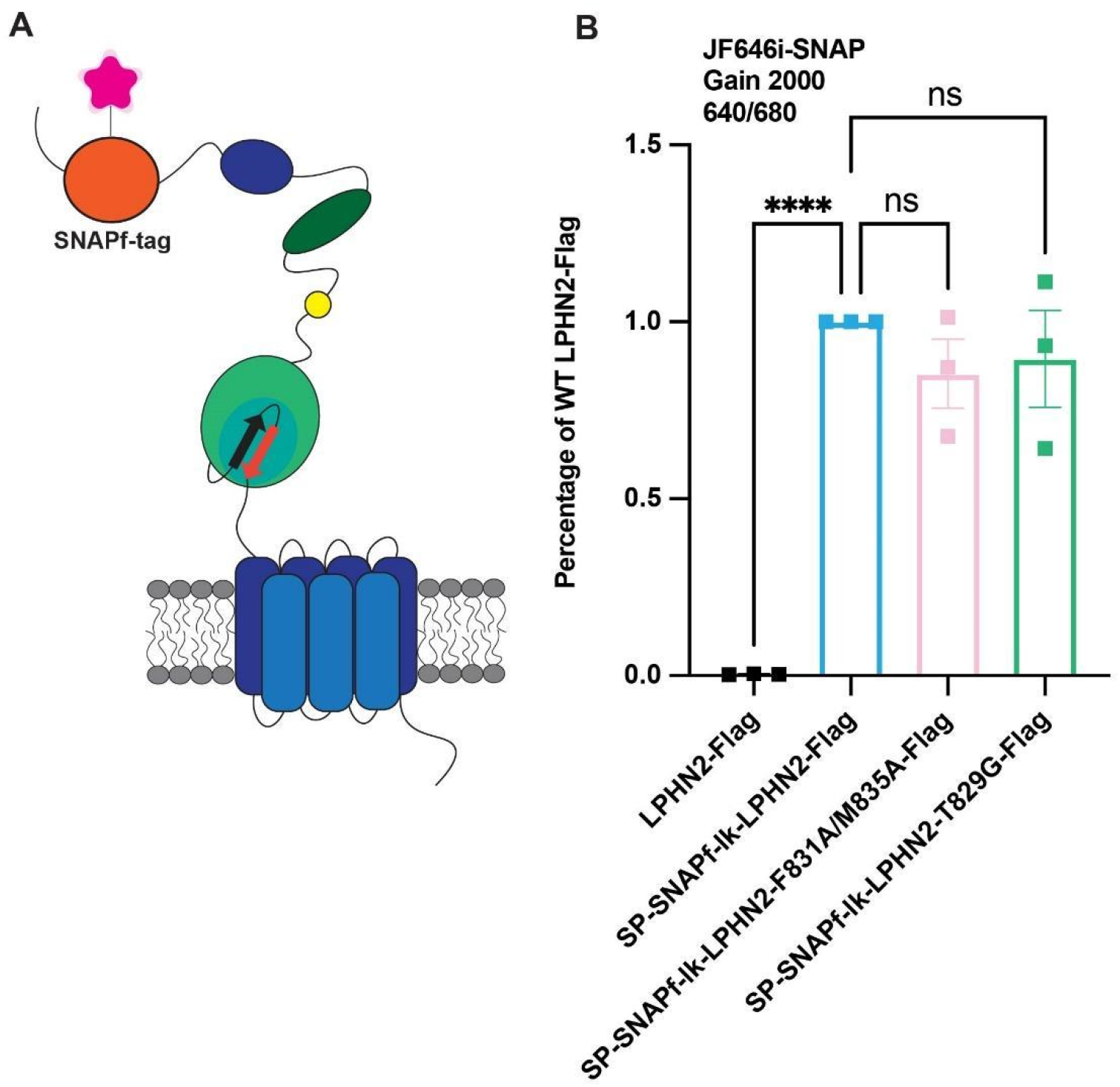
Lphn2, Lphn2-F831A/M835A, and Lphn2-T829G are expressed at the cell surface at comparable levels. **(A)** Schematic of Lphn2 constructs with an additional SNAPfast-tag on the N terminus. **(B)** Lphn2 constructs tagged with a SNAPfast-tag were expressed in HEK293T cells for 24 hr. Transfected cells were incubated for 30 min with 1 µM membrane-impermeant Janelia Fluor 646, a SNAP-tag substrate. Emission was read using the filter 640/680 at a Gain 2000 using a PHERAstar FS microplate reader. Expression was calculated as a percentage of WT Lphn2. Statistical analysis was conducted using a one-way ANOVA followed by Dunnett’s multiple comparisons test (****, <0.0001).

**Figure 3—figure supplement 2.**
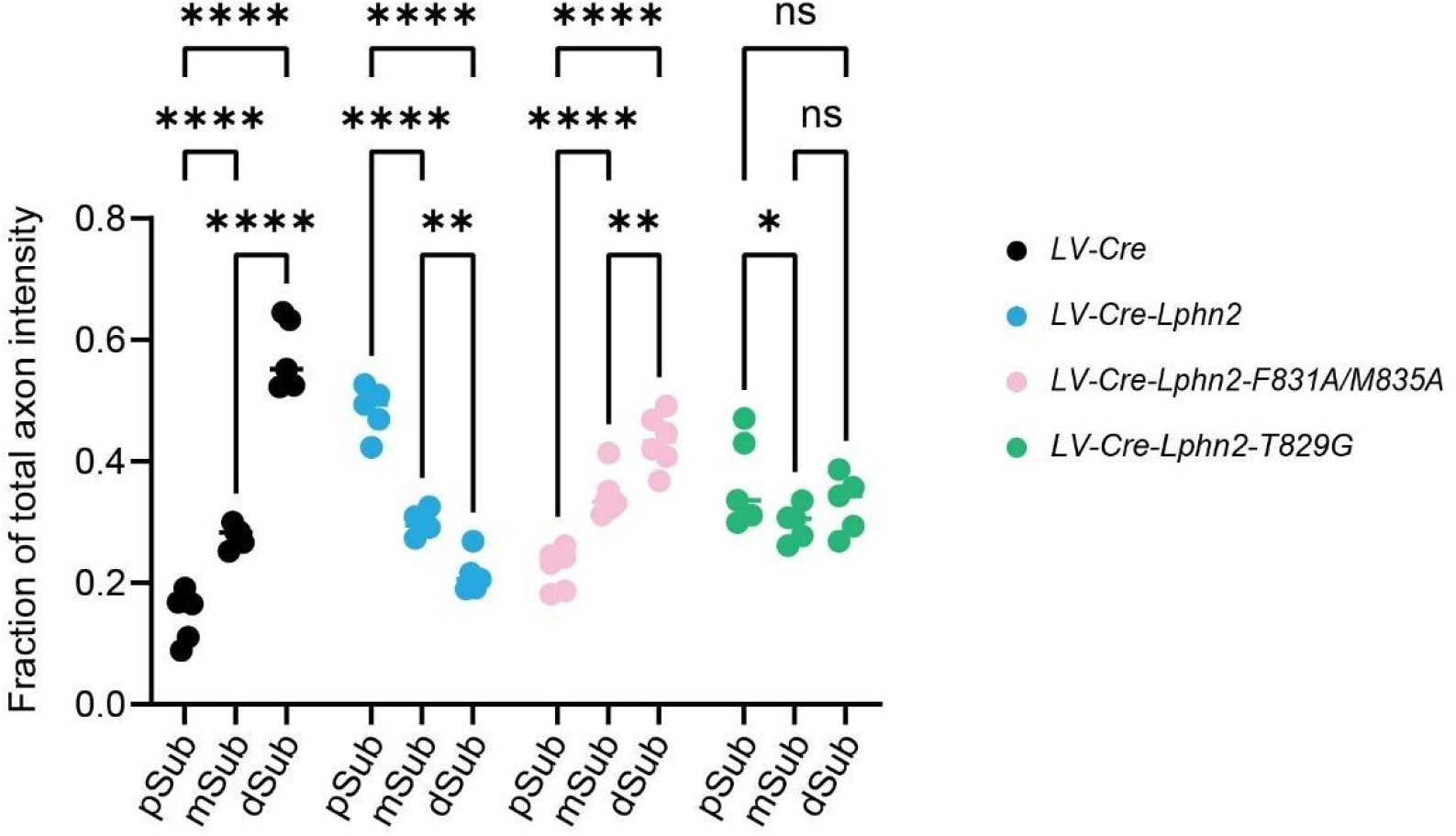
Comparison of fraction of total axon intensity across the subiculum within the same experimental condition. *LV-Cre* animals have low axon intensity in pSub and high axon intensity in dSub. LV-Cre-Lphn2 animals have high axon intensity in pSub and low axon intensity in dSub. LV-Cre-Lphn2-F831A/M835A animals show a similar pattern to LV-Cre with increasing axon intensity from pSub to dSub. LV-Cre-Lphn2-T829G shows moderate axon enrichment in pSub but overall the total axon intensity is not biased towards any one region of subiculum. Means ± SEM; two-way ANOVA with Sidak’s multiple comparisons test.

**Figure 5—figure supplement 1.**
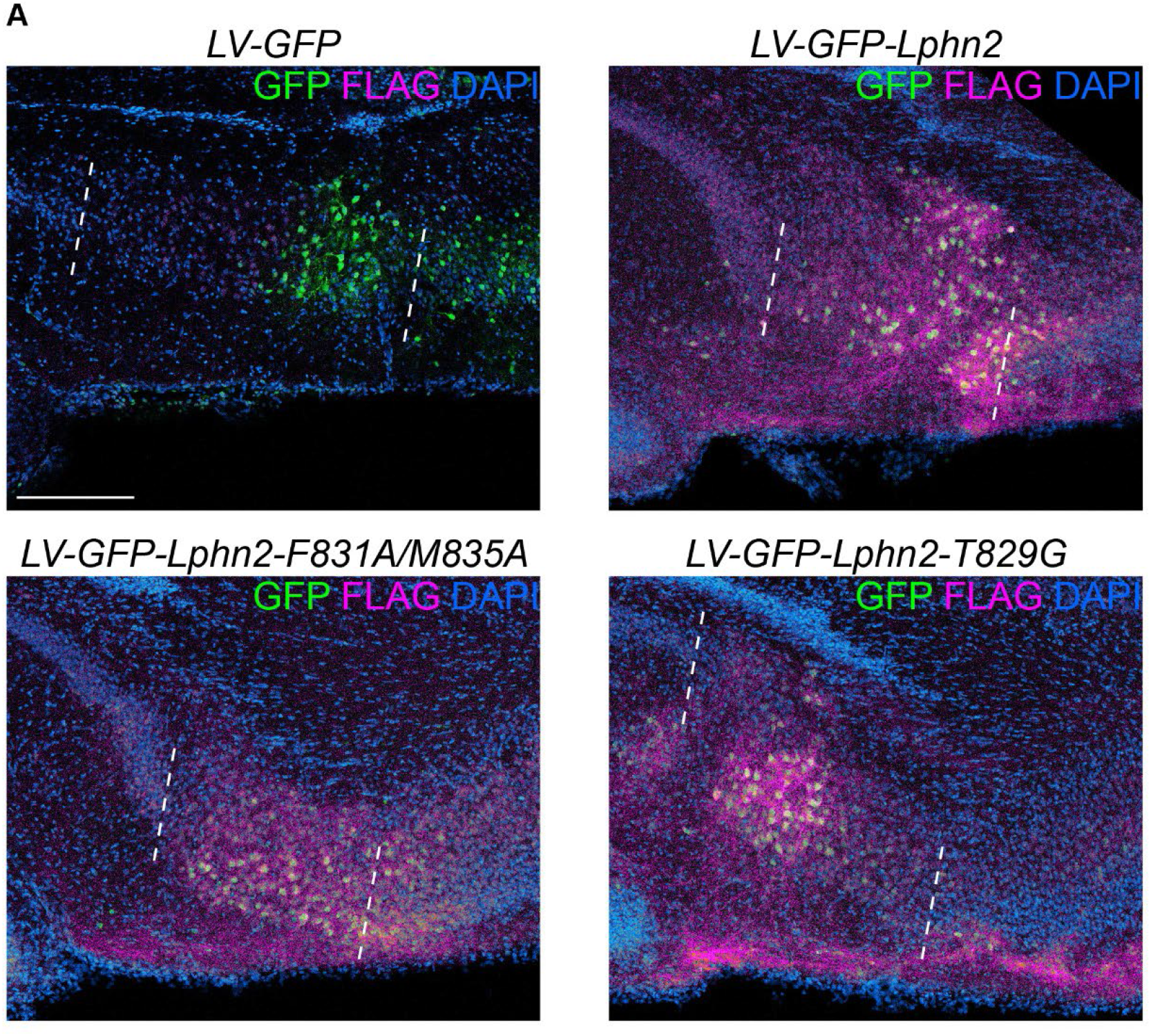
*In vivo* expression of lentivirus used in Figure 5. (**A**) Representative images of GFP and FLAG immunostaining in P8 subiculum of mice injected with lentiviruses used in Figure 5. The region between the white dashed line is subiculum. Scale bars represents 200 μm.

**Figure 2 – Source data.**
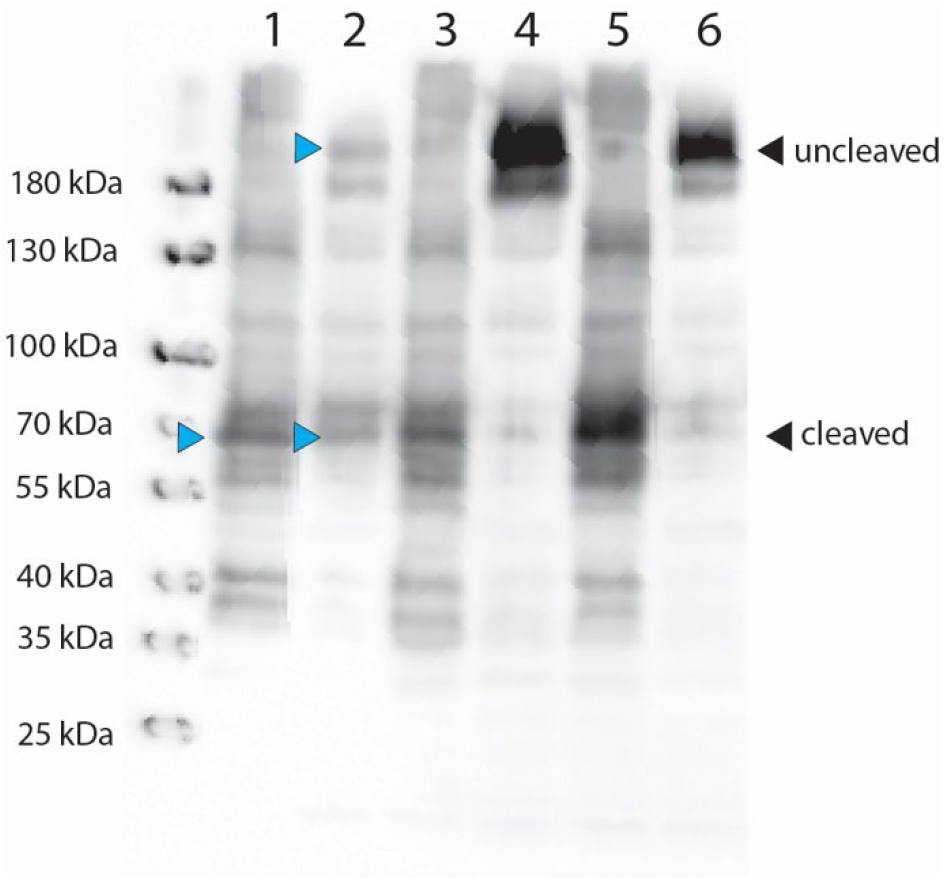
Uncropped immunoblot analysis of Lphn2 expression in HEK293T cells. Primary antibody against Flag (1:500, ThermoFisher, PA1-984B), secondary antibody anti-rabbit HRP (1:10,000, ThermoFisher, Cat #31458). Blue arrows indicate bands of interest for 1: Lphn2-CTF-Flag and 2: Full-length Lphn2-Flag.

**Figure 3 – Source data.**
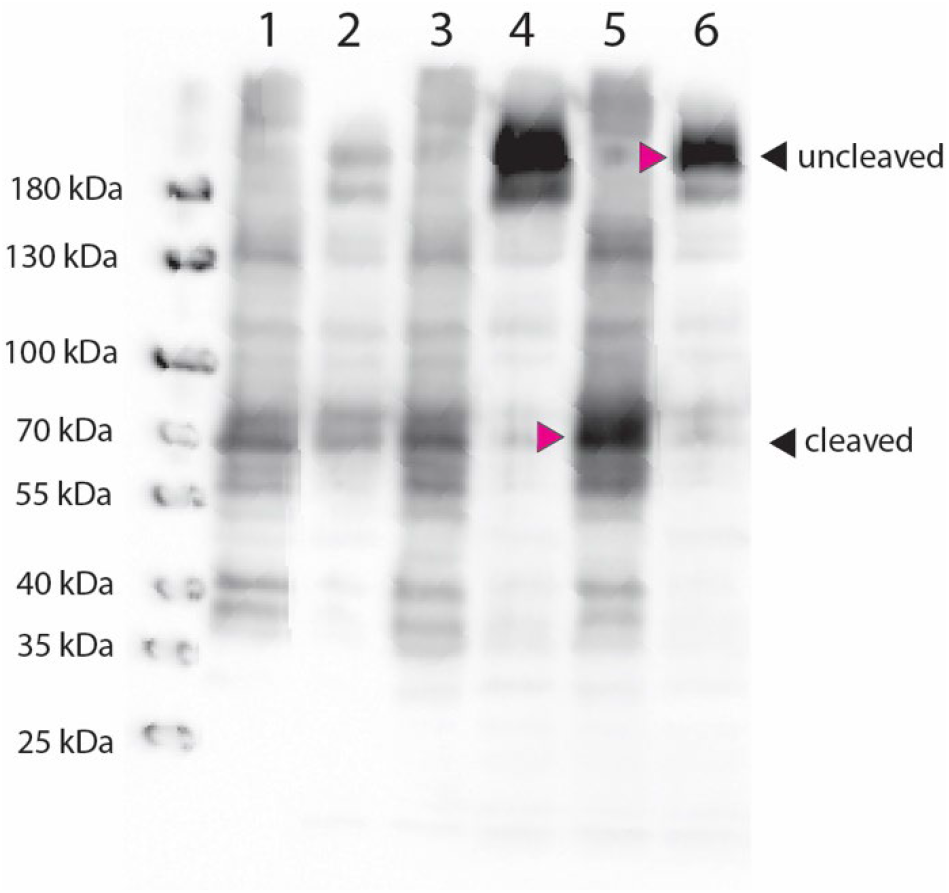
Uncropped immunoblot analysis of Lphn2 expression in HEK293T cells. Primary antibody against Flag (1:500, ThermoFisher, PA1-984B), secondary antibody anti-rabbit HRP (1:10,000, ThermoFisher, Cat #31458). Magenta arrows indicate bands of interest for 5: Lphn2-F831A/M835A-CTF-Flag and 6: Full-length Lphn2-F831A/M835A-Flag.

**Figure 4 – Source data.**
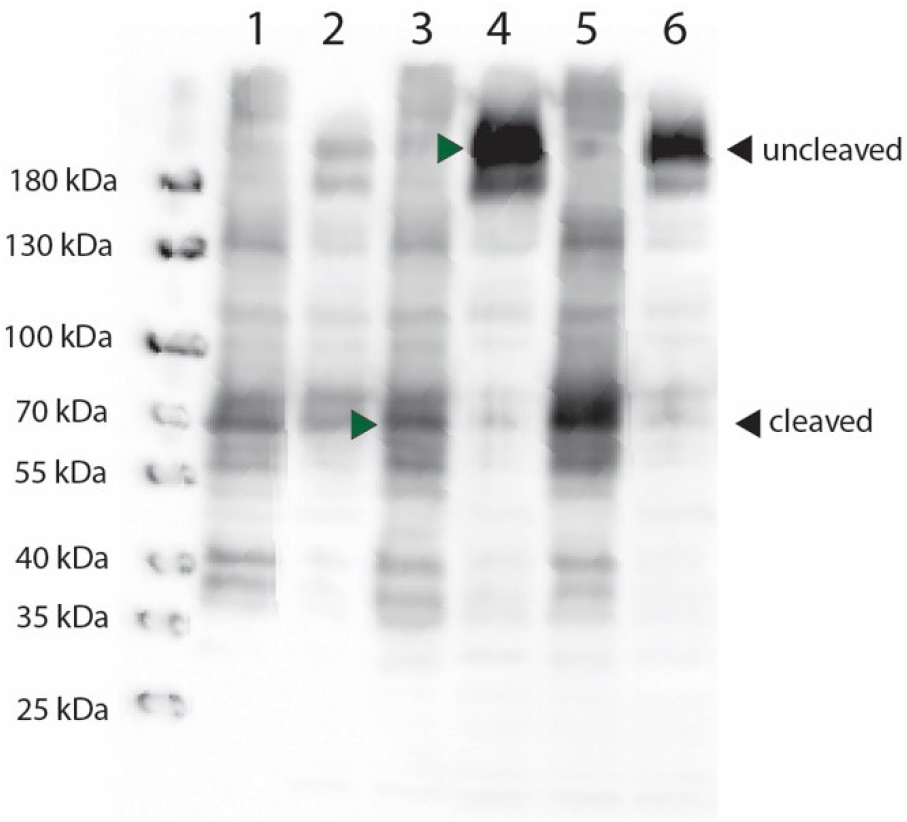
Uncropped immunoblot analysis of Lphn2 expression in HEK293T cells. Primary antibody against Flag (1:500, ThermoFisher, PA1-984B), secondary antibody anti-rabbit HRP (1:10,000, ThermoFisher, Cat #31458). Green arrows indicate bands of interest for 3: Lphn2-T829G-CTF-Flag and 4: Full-length Lphn2-T829G-Flag.

## Notes

### Competing Interest Statement

The authors have declared no competing interest.

